## Supplementary material for "A systematic evaluation of highly variable gene selection methods for single-cell RNA-sequencing": The full latex folder for manuscript: sn-article.bib

% bmc_article.bib
% 
%  An example of bibtex entries.
%  Entries taken from BMC instructions for authors page.

% uncomment next line to make author-year bibliography
% @settings{label, options="nameyear"}
@article{lee2020single,
  title={Single-cell multiomics: technologies and data analysis methods},
  author={Lee, Jeongwoo and Hyeon, Do Young and Hwang, Daehee},
  journal={Experimental \& Molecular Medicine},
  volume={52},
  number={9},
  pages={1428--1442},
  year={2020},
  publisher={Nature Publishing Group}
}

@article{haque2017practical,
  title={A practical guide to single-cell RNA-sequencing for biomedical research and clinical applications},
  author={Haque, Ashraful and Engel, Jessica and Teichmann, Sarah A and L{\"o}nnberg, Tapio},
  journal={Genome medicine},
  volume={9},
  number={1},
  pages={1--12},
  year={2017},
  publisher={BioMed Central}
}

@article{satija2015spatial,
  title={Spatial reconstruction of single-cell gene expression data},
  author={Satija, Rahul and Farrell, Jeffrey A and Gennert, David and Schier, Alexander F and Regev, Aviv},
  journal={Nature biotechnology},
  volume={33},
  number={5},
  pages={495--502},
  year={2015},
  publisher={Nature Publishing Group}
}


@article{yip2019evaluation,
  title={Evaluation of tools for highly variable gene discovery from single-cell RNA-seq data},
  author={Yip, Shun H and Sham, Pak Chung and Wang, Junwen},
  journal={Briefings in bioinformatics},
  volume={20},
  number={4},
  pages={1583--1589},
  year={2019},
  publisher={Oxford University Press}
}

@article{townes2019feature,
  title={Feature selection and dimension reduction for single-cell RNA-Seq based on a multinomial model},
  author={Townes, F William and Hicks, Stephanie C and Aryee, Martin J and Irizarry, Rafael A},
  journal={Genome biology},
  volume={20},
  number={1},
  pages={1--16},
  year={2019},
  publisher={BioMed Central}
}


@article{lun2016step,
  title={A step-by-step workflow for low-level analysis of single-cell RNA-seq data with Bioconductor},
  author={Lun, Aaron TL and McCarthy, Davis J and Marioni, John C},
  journal={F1000Research},
  volume={5},
  year={2016},
  publisher={Faculty of 1000 Ltd}
}


@article{hafemeister2019normalization,
  title={Normalization and variance stabilization of single-cell RNA-seq data using regularized negative binomial regression},
  author={Hafemeister, Christoph and Satija, Rahul},
  journal={Genome biology},
  volume={20},
  number={1},
  pages={1--15},
  year={2019},
  publisher={Springer}
}

@article{wolf2018scanpy,
  title={SCANPY: large-scale single-cell gene expression data analysis},
  author={Wolf, F Alexander and Angerer, Philipp and Theis, Fabian J},
  journal={Genome biology},
  volume={19},
  number={1},
  pages={1--5},
  year={2018},
  publisher={BioMed Central}
}


@article{hou2020systematic,
  title={A systematic evaluation of single-cell RNA-sequencing imputation methods},
  author={Hou, Wenpin and Ji, Zhicheng and Ji, Hongkai and Hicks, Stephanie C},
  journal={Genome biology},
  volume={21},
  number={1},
  pages={1--30},
  year={2020},
  publisher={Springer}
}

@article{booeshaghi2022depth,
  title={Depth normalization for single-cell genomics count data},
  author={Booeshaghi, A Sina and Hallgr{\'\i}msd{\'o}ttir, Ingileif B and G{\'a}lvez-Merch{\'a}n, {\'A}ngel and Pachter, Lior},
  journal={bioRxiv},
  year={2022},
  publisher={Cold Spring Harbor Laboratory}
}

@article{mcinnes2018umap,
  title={Umap: Uniform manifold approximation and projection for dimension reduction},
  author={McInnes, Leland and Healy, John and Melville, James},
  journal={arXiv preprint arXiv:1802.03426},
  year={2018}
}


@article{stuart2019comprehensive,
  title={Comprehensive integration of single-cell data},
  author={Stuart, Tim and Butler, Andrew and Hoffman, Paul and Hafemeister, Christoph and Papalexi, Efthymia and Mauck III, William M and Hao, Yuhan and Stoeckius, Marlon and Smibert, Peter and Satija, Rahul},
  journal={Cell},
  volume={177},
  number={7},
  pages={1888--1902},
  year={2019},
  publisher={Elsevier}
}


@inproceedings{santos2009use,
  title={On the use of the adjusted rand index as a metric for evaluating supervised classification},
  author={Santos, Jorge M and Embrechts, Mark},
  booktitle={International conference on artificial neural networks},
  pages={175--184},
  year={2009},
  organization={Springer}
}


@article{han2018mapping,
  title={Mapping the mouse cell atlas by microwell-seq},
  author={Han, Xiaoping and Wang, Renying and Zhou, Yincong and Fei, Lijiang and Sun, Huiyu and Lai, Shujing and Saadatpour, Assieh and Zhou, Ziming and Chen, Haide and Ye, Fang and others},
  journal={Cell},
  volume={172},
  number={5},
  pages={1091--1107},
  year={2018},
  publisher={Elsevier}
}


@article{picelli2014full,
  title={Full-length RNA-seq from single cells using Smart-seq2},
  author={Picelli, Simone and Faridani, Omid R and Bj{\"o}rklund, {\AA}sa K and Winberg, G{\"o}sta and Sagasser, Sven and Sandberg, Rickard},
  journal={Nature protocols},
  volume={9},
  number={1},
  pages={171--181},
  year={2014},
  publisher={Nature Publishing Group}
}

@article{zheng2017massively,
  title={Massively parallel digital transcriptional profiling of single cells},
  author={Zheng, Grace XY and Terry, Jessica M and Belgrader, Phillip and Ryvkin, Paul and Bent, Zachary W and Wilson, Ryan and Ziraldo, Solongo B and Wheeler, Tobias D and McDermott, Geoff P and Zhu, Junjie and others},
  journal={Nature communications},
  volume={8},
  number={1},
  pages={1--12},
  year={2017},
  publisher={Nature Publishing Group}
}


@article{darmanis2017single,
  title={Single-cell RNA-seq analysis of infiltrating neoplastic cells at the migrating front of human glioblastoma},
  author={Darmanis, Spyros and Sloan, Steven A and Croote, Derek and Mignardi, Marco and Chernikova, Sophia and Samghababi, Peyman and Zhang, Ye and Neff, Norma and Kowarsky, Mark and Caneda, Christine and others},
  journal={Cell reports},
  volume={21},
  number={5},
  pages={1399--1410},
  year={2017},
  publisher={Elsevier}
}


@article{duo2018systematic,
  title={A systematic performance evaluation of clustering methods for single-cell RNA-seq data},
  author={Du{\`o}, Angelo and Robinson, Mark D and Soneson, Charlotte},
  journal={F1000Research},
  volume={7},
  year={2018},
  publisher={Faculty of 1000 Ltd}
}


@article{linderman2022zero,
  title={Zero-preserving imputation of single-cell RNA-seq data},
  author={Linderman, George C and Zhao, Jun and Roulis, Manolis and Bielecki, Piotr and Flavell, Richard A and Nadler, Boaz and Kluger, Yuval},
  journal={Nature communications},
  volume={13},
  number={1},
  pages={1--11},
  year={2022},
  publisher={Nature Publishing Group}
}


@article{stoeckius2017large,
  title={Large-scale simultaneous measurement of epitopes and transcriptomes in single cells},
  author={Stoeckius, Marlon and Hafemeister, Christoph and Stephenson, William and Houck-Loomis, Brian and Chattopadhyay, Pratip K and Swerdlow, Harold and Satija, Rahul and Smibert, Peter},
  journal={Nature methods},
  volume={14},
  number={9},
  pages={865},
  year={2017},
  publisher={NIH Public Access}
}


@article{jardine2021blood,
  title={Blood and immune development in human fetal bone marrow and Down syndrome},
  author={Jardine, Laura and Webb, Simone and Goh, Issac and Quiroga Londo{\~n}o, Mariana and Reynolds, Gary and Mather, Michael and Olabi, Bayanne and Stephenson, Emily and Botting, Rachel A and Horsfall, Dave and others},
  journal={Nature},
  volume={598},
  number={7880},
  pages={327--331},
  year={2021},
  publisher={Nature Publishing Group}
}


@article{hao2021integrated,
  title={Integrated analysis of multimodal single-cell data},
  author={Hao, Yuhan and Hao, Stephanie and Andersen-Nissen, Erica and Mauck III, William M and Zheng, Shiwei and Butler, Andrew and Lee, Maddie J and Wilk, Aaron J and Darby, Charlotte and Zager, Michael and others},
  journal={Cell},
  volume={184},
  number={13},
  pages={3573--3587},
  year={2021},
  publisher={Elsevier}
}

@article{su2020multi,
  title={Multi-omics resolves a sharp disease-state shift between mild and moderate COVID-19},
  author={Su, Yapeng and Chen, Daniel and Yuan, Dan and Lausted, Christopher and Choi, Jongchan and Dai, Chengzhen L and Voillet, Valentin and Duvvuri, Venkata R and Scherler, Kelsey and Troisch, Pamela and others},
  journal={Cell},
  volume={183},
  number={6},
  pages={1479--1495},
  year={2020},
  publisher={Elsevier}
}


@article{chen2019high,
  title={High-throughput sequencing of the transcriptome and chromatin accessibility in the same cell},
  author={Chen, Song and Lake, Blue B and Zhang, Kun},
  journal={Nature biotechnology},
  volume={37},
  number={12},
  pages={1452--1457},
  year={2019},
  publisher={Nature Publishing Group}
}

@article{rank_vogt1999fusion,
  title={Fusion via a linear combination of scores},
  author={Vogt, Christopher C and Cottrell, Garrison W},
  journal={Information retrieval},
  volume={1},
  number={3},
  pages={151--173},
  year={1999},
  publisher={Springer}
}

@inproceedings{rank_aslam2001models,
  title={Models for metasearch},
  author={Aslam, Javed A and Montague, Mark},
  booktitle={Proceedings of the 24th annual international ACM SIGIR conference on Research and development in information retrieval},
  pages={276--284},
  year={2001}
}

@inproceedings{rank_fagin2003efficient,
  title={Efficient similarity search and classification via rank aggregation},
  author={Fagin, Ronald and Kumar, Ravi and Sivakumar, Dandapani},
  booktitle={Proceedings of the 2003 ACM SIGMOD international conference on Management of data},
  pages={301--312},
  year={2003}
}

@article{tang2009mrna,
  title={mRNA-Seq whole-transcriptome analysis of a single cell},
  author={Tang, Fuchou and Barbacioru, Catalin and Wang, Yangzhou and Nordman, Ellen and Lee, Clarence and Xu, Nanlan and Wang, Xiaohui and Bodeau, John and Tuch, Brian B and Siddiqui, Asim and others},
  journal={Nature methods},
  volume={6},
  number={5},
  pages={377--382},
  year={2009},
  publisher={Nature Publishing Group US New York}
}


@article{miao2020putative,
  title={Putative cell type discovery from single-cell gene expression data},
  author={Miao, Zhichao and Moreno, Pablo and Huang, Ni and Papatheodorou, Irene and Brazma, Alvis and Teichmann, Sarah A},
  journal={Nature methods},
  volume={17},
  number={6},
  pages={621--628},
  year={2020},
  publisher={Nature Publishing Group US New York}
}

@article{kotliar2019identifying,
  title={Identifying gene expression programs of cell-type identity and cellular activity with single-cell RNA-Seq},
  author={Kotliar, Dylan and Veres, Adrian and Nagy, M Aurel and Tabrizi, Shervin and Hodis, Eran and Melton, Douglas A and Sabeti, Pardis C},
  journal={Elife},
  volume={8},
  pages={e43803},
  year={2019},
  publisher={eLife Sciences Publications, Ltd}
}

@article{zheng2017landscape,
  title={Landscape of infiltrating T cells in liver cancer revealed by single-cell sequencing},
  author={Zheng, Chunhong and Zheng, Liangtao and Yoo, Jae-Kwang and Guo, Huahu and Zhang, Yuanyuan and Guo, Xinyi and Kang, Boxi and Hu, Ruozhen and Huang, Julie Y and Zhang, Qiming and others},
  journal={Cell},
  volume={169},
  number={7},
  pages={1342--1356},
  year={2017},
  publisher={Elsevier}
}

@Manual{mixmix2024,
  title = {mixhvg: Mixture of Multiple Highly Variable Feature Selection Methods},
  author = {Ruzhang Zhao},
  year = {2023},
  note = {R package version 0.1.1},
  url = {https://CRAN.R-project.org/package=mixhvg}
}


@article{kriegeskorte2009circular,
  title={Circular analysis in systems neuroscience: the dangers of double dipping},
  author={Kriegeskorte, Nikolaus and Simmons, W Kyle and Bellgowan, Patrick SF and Baker, Chris I},
  journal={Nature neuroscience},
  volume={12},
  number={5},
  pages={535--540},
  year={2009},
  publisher={Nature Publishing Group US New York}
}
@article{taylor2015statistical,
  title={Statistical learning and selective inference},
  author={Taylor, Jonathan and Tibshirani, Robert J},
  journal={Proceedings of the National Academy of Sciences},
  volume={112},
  number={25},
  pages={7629--7634},
  year={2015},
  publisher={National Acad Sciences}
}

@article{ji2016tscan,
  title={TSCAN: Pseudo-time reconstruction and evaluation in single-cell RNA-seq analysis},
  author={Ji, Zhicheng and Ji, Hongkai},
  journal={Nucleic acids research},
  volume={44},
  number={13},
  pages={e117--e117},
  year={2016},
  publisher={Oxford University Press}
}


@article{wang2019data,
  title={Data denoising with transfer learning in single-cell transcriptomics},
  author={Wang, Jingshu and Agarwal, Divyansh and Huang, Mo and Hu, Gang and Zhou, Zilu and Ye, Chengzhong and Zhang, Nancy R},
  journal={Nature methods},
  volume={16},
  number={9},
  pages={875--878},
  year={2019},
  publisher={Nature Publishing Group US New York}
}

@article{cang2023screening,
  title={Screening cell--cell communication in spatial transcriptomics via collective optimal transport},
  author={Cang, Zixuan and Zhao, Yanxiang and Almet, Axel A and Stabell, Adam and Ramos, Raul and Plikus, Maksim V and Atwood, Scott X and Nie, Qing},
  journal={Nature Methods},
  volume={20},
  number={2},
  pages={218--228},
  year={2023},
  publisher={Nature Publishing Group US New York}
}

@article{wold1987principal,
  title={Principal component analysis},
  author={Wold, Svante and Esbensen, Kim and Geladi, Paul},
  journal={Chemometrics and intelligent laboratory systems},
  volume={2},
  number={1-3},
  pages={37--52},
  year={1987},
  publisher={Elsevier}
}

@article{blondel2008fast,
  title={Fast unfolding of communities in large networks},
  author={Blondel, Vincent D and Guillaume, Jean-Loup and Lambiotte, Renaud and Lefebvre, Etienne},
  journal={Journal of statistical mechanics: theory and experiment},
  volume={2008},
  number={10},
  pages={P10008},
  year={2008},
  publisher={IOP Publishing}
}

@article{deerwester1990indexing,
  title={Indexing by latent semantic analysis},
  author={Deerwester, Scott and Dumais, Susan T and Furnas, George W and Landauer, Thomas K and Harshman, Richard},
  journal={Journal of the American society for information science},
  volume={41},
  number={6},
  pages={391--407},
  year={1990},
  publisher={Wiley Online Library}
}
@article{trapnell2014dynamics,
  title={The dynamics and regulators of cell fate decisions are revealed by pseudotemporal ordering of single cells},
  author={Trapnell, Cole and Cacchiarelli, Davide and Grimsby, Jonna and Pokharel, Prapti and Li, Shuqiang and Morse, Michael and Lennon, Niall J and Livak, Kenneth J and Mikkelsen, Tarjei S and Rinn, John L},
  journal={Nature biotechnology},
  volume={32},
  number={4},
  pages={381--386},
  year={2014},
  publisher={Nature Publishing Group US New York}
}

@article{stuart2021single,
  title={Single-cell chromatin state analysis with Signac},
  author={Stuart, Tim and Srivastava, Avi and Madad, Shaista and Lareau, Caleb A and Satija, Rahul},
  journal={Nature methods},
  volume={18},
  number={11},
  pages={1333--1341},
  year={2021},
  publisher={Nature Publishing Group US New York}
}


@article{butler2018integrating,
  title={Integrating single-cell transcriptomic data across different conditions, technologies, and species},
  author={Butler, Andrew and Hoffman, Paul and Smibert, Peter and Papalexi, Efthymia and Satija, Rahul},
  journal={Nature biotechnology},
  volume={36},
  number={5},
  pages={411--420},
  year={2018},
  publisher={Nature Publishing Group US New York}
}


@article{lancichinetti2009detecting,
  title={Detecting the overlapping and hierarchical community structure in complex networks},
  author={Lancichinetti, Andrea and Fortunato, Santo and Kert{\'e}sz, J{\'a}nos},
  journal={New journal of physics},
  volume={11},
  number={3},
  pages={033015},
  year={2009},
  publisher={IOP Publishing}
}


@article{rousseeuw1987silhouettes,
  title={Silhouettes: a graphical aid to the interpretation and validation of cluster analysis},
  author={Rousseeuw, Peter J},
  journal={Journal of computational and applied mathematics},
  volume={20},
  pages={53--65},
  year={1987},
  publisher={Elsevier}
}

@article{luecken2022benchmarking,
  title={Benchmarking atlas-level data integration in single-cell genomics},
  author={Luecken, Malte D and B{\"u}ttner, Maren and Chaichoompu, Kridsadakorn and Danese, Anna and Interlandi, Marta and M{\"u}ller, Michaela F and Strobl, Daniel C and Zappia, Luke and Dugas, Martin and Colom{\'e}-Tatch{\'e}, Maria and others},
  journal={Nature methods},
  volume={19},
  number={1},
  pages={41--50},
  year={2022},
  publisher={Nature Publishing Group US New York}
}


@article{calinski1974dendrite,
  title={A dendrite method for cluster analysis},
  author={Cali{\'n}ski, Tadeusz and Harabasz, Jerzy},
  journal={Communications in Statistics-theory and Methods},
  volume={3},
  number={1},
  pages={1--27},
  year={1974},
  publisher={Taylor \& Francis}
}

@article{zhang2021tree,
  title={Tree-based Correlation Screen and Visualization for Exploring Phenotype-Cell Type Association in Multiple Sample Single-Cell RNA-Sequencing Experiments},
  author={Zhang, Boyang and Ji, Zhicheng and Ji, Hongkai},
  journal={bioRxiv},
  pages={2021--10},
  year={2021},
  publisher={Cold Spring Harbor Laboratory}
}


@article{korsunsky2019fast,
  title={Fast, sensitive and accurate integration of single-cell data with Harmony},
  author={Korsunsky, Ilya and Millard, Nghia and Fan, Jean and Slowikowski, Kamil and Zhang, Fan and Wei, Kevin and Baglaenko, Yuriy and Brenner, Michael and Loh, Po-ru and Raychaudhuri, Soumya},
  journal={Nature methods},
  volume={16},
  number={12},
  pages={1289--1296},
  year={2019},
  publisher={Nature Publishing Group US New York}
}
