## Supplementary material for "A systematic evaluation of highly variable gene selection methods for single-cell RNA-sequencing": The full latex folder for manuscript: sn-article.tex

%Version 3 December 2023
% See section 11 of the User Manual for version history
%
%%%%%%%%%%%%%%%%%%%%%%%%%%%%%%%%%%%%%%%%%%%%%%%%%%%%%%%%%%%%%%%%%%%%%%
%%                                                                 %%
%% Please do not use \input{...} to include other tex files.       %%
%% Submit your LaTeX manuscript as one .tex document.              %%
%%                                                                 %%
%% All additional figures and files should be attached             %%
%% separately and not embedded in the \TeX\ document itself.       %%
%%                                                                 %%
%%%%%%%%%%%%%%%%%%%%%%%%%%%%%%%%%%%%%%%%%%%%%%%%%%%%%%%%%%%%%%%%%%%%%

%%\documentclass[referee,sn-basic]{sn-jnl}% referee option is meant for double line spacing

%%=======================================================%%
%% to print line numbers in the margin use lineno option %%
%%=======================================================%%

%%\documentclass[lineno,sn-basic]{sn-jnl}% Basic Springer Nature Reference Style/Chemistry Reference Style

%%======================================================%%
%% to compile with pdflatex/xelatex use pdflatex option %%
%%======================================================%%

%%\documentclass[pdflatex,sn-basic]{sn-jnl}% Basic Springer Nature Reference Style/Chemistry Reference Style


%%Note: the following reference styles support Namedate and Numbered referencing. By default the style follows the most common style. To switch between the options you can add or remove  in the optional parenthesis. 
%%The option is available for: sn-basic.bst, sn-vancouver.bst, sn-chicago.bst%  
 
%%\documentclass[pdflatex,sn-nature]{sn-jnl}% Style for submissions to Nature Portfolio journals
%%\documentclass[pdflatex,sn-basic]{sn-jnl}% Basic Springer Nature Reference Style/Chemistry Reference Style
\documentclass[pdflatex,sn-mathphys-num]{sn-jnl}% Math and Physical Sciences Numbered Reference Style 
%%\documentclass[pdflatex,sn-mathphys-ay]{sn-jnl}% Math and Physical Sciences Author Year Reference Style
%%\documentclass[pdflatex,sn-aps]{sn-jnl}% American Physical Society (APS) Reference Style
%%\documentclass[pdflatex,sn-vancouver,Numbered]{sn-jnl}% Vancouver Reference Style
%%\documentclass[pdflatex,sn-apa]{sn-jnl}% APA Reference Style 
%%\documentclass[pdflatex,sn-chicago]{sn-jnl}% Chicago-based Humanities Reference Style

%%%% Standard Packages
%%<additional latex packages if required can be included here>

\usepackage{graphicx}%
\usepackage{multirow}%
\usepackage{amsmath,amssymb,amsfonts}%
\usepackage{amsthm}%
\usepackage{mathrsfs}%
\usepackage[title]{appendix}%
\usepackage{xcolor}%
\usepackage{textcomp}%
\usepackage{manyfoot}%
%%%%
\usepackage{hyperref}
%\usepackage[export]{adjustbox}
\usepackage{tabularx}

%%%%%=============================================================================%%%%
%%%%  Remarks: This template is provided to aid authors with the preparation
%%%%  of original research articles intended for submission to journals published 
%%%%  by Springer Nature. The guidance has been prepared in partnership with 
%%%%  production teams to conform to Springer Nature technical requirements. 
%%%%  Editorial and presentation requirements differ among journal portfolios and 
%%%%  research disciplines. You may find sections in this template are irrelevant 
%%%%  to your work and are empowered to omit any such section if allowed by the 
%%%%  journal you intend to submit to. The submission guidelines and policies 
%%%%  of the journal take precedence. A detailed User Manual is available in the 
%%%%  template package for technical guidance.
%%%%%=============================================================================%%%%

%% as per the requirement new theorem styles can be included as shown below  meant for continuous numbers
%%\newtheorem{theorem}{Theorem}[section]% meant for sectionwise numbers
%% optional argument [theorem] produces theorem numbering sequence instead of independent numbers for Proposition
%\newtheorem{proposition}[theorem]{Proposition}% 
%%\newtheorem{proposition}{Proposition}% to get separate numbers for theorem and proposition etc.

%\theoremstyle{thmstyletwo}%
%\newtheorem{example}{Example}%
%\newtheorem{remark}{Remark}%

%\theoremstyle{thmstylethree}%
%\newtheorem{definition}{Definition}%

\raggedbottom
%%\unnumbered% uncomment this for unnumbered level heads

\begin{document}

\title[HVG selection]{A systematic evaluation of highly variable gene selection methods for single-cell RNA-sequencing}

%%=============================================================%%
%% GivenName	-> \fnm{Joergen W.}
%% Particle	-> \spfx{van der} -> surname prefix
%% FamilyName	-> \sur{Ploeg}
%% Suffix	-> \sfx{IV}
%% \author*[1,2]{\fnm{Joergen W.} \spfx{van der} \sur{Ploeg} 
%%=============================================================%%

\author[1]{\fnm{Jiuyao} \sur{Lu}}
\author[1]{\fnm{Weiqiang} \sur{Zhou}}
\author[1]{\fnm{Ni} \sur{Zhao}}

\affil*[1]{\orgdiv{Department of Biostatistics}, \orgname{Johns Hopkins Bloomberg School of Public Health}, \orgaddress{ \city{Baltimore}, \postcode{21212}, \state{MD}, \country{USA}}}
%%==================================%%
%% Sample for unstructured abstract %%
%%==================================%%

\abstract{

\textbf{Background}:
Selecting highly variable features is a crucial step in most analysis pipelines of single-cell RNA-sequencing (scRNA-seq) data. Despite numerous methods proposed in recent years, a systematic understanding of the best solution is still lacking. 

\textbf{Results}: 
Here, we systematically evaluate 47 highly variable gene (HVG) selection methods, consisting of 21 baseline methods developed based on different data transformations and mean-variance adjustment techniques and 26 hybrid methods developed based on mixtures of baseline methods. Across 19 diverse benchmark datasets, 18 objective evaluation criteria per method, and 5,358 analysis settings, we observe that no single baseline method consistently outperforms the others across all datasets and criteria. However, hybrid methods as a group robustly outperform individual baseline methods. Based on these findings, a new HVG selection approach, mixture HVG selection (mixHVG), that incorporates top-ranked features from multiple baseline methods is proposed as a better solution to HVG selection. An open source R package \texttt{mixhvg} is developed to enable convenient use of mixHVG and its integration into users' data analysis pipelines. 

\textbf{Conclusion}: 
Our benchmark study not only provides a systematic comparison of existing methods, leading to a better HVG selection solution, but also creates a pipeline and resource consisting of diverse benchmark data and criteria for evaluating new methods in the future. 
}
\keywords{Single-cell Genomics, Preprocessing, Feature Selection, Benchmark, Software}
\maketitle

\section*{Background}
Single-cell RNA-sequencing (scRNA-seq) has been rapidly transforming biomedical research \cite{tang2009mrna,haque2017practical}. 
By allowing gene expression in individual cells to be examined, this technology facilitates the comprehensive assessment of the gene expression landscape within diverse cell populations.
Its applications span from the identification of novel cell types \cite{stuart2019comprehensive,miao2020putative,kotliar2019identifying} to the reconstruction of cells' temporal and spatial transcriptional profiles \cite{zheng2017landscape}, and the investigation of cell-cell interactions and communication \cite{cang2023screening}. 
However, scRNA-seq data are inherently high-dimensional and noisy. Consequently, effective data analysis is pivotal for distilling valuable insights from raw data and uncovering novel discoveries.

The selection of highly variable genes (HVGs), referred to interchangeably as highly variable features (HVFs) in this article, is a crucial step in many scRNA-seq data analysis pipelines, influencing the majority of subsequent analytical tasks. For instance, in a standard data analysis pipeline, Seurat \cite{satija2015spatial}, following data quality control and normalization, genes demonstrating the greatest variation across cells (i.e., HVGs) are identified. Subsequent data embedding such as principal component analysis (PCA)  \cite{wold1987principal}, visualization such as Uniform Manifold Approximation and Projection (UMAP) \cite{mcinnes2018umap}, and downstream analyses including cell clustering \cite{blondel2008fast} and pseudotemporal trajectory inference \cite{trapnell2014dynamics}, all rely on the selected HVGs. The primary objective of HVG selection is to address the curse of dimensionality inherent in high-dimensional data by filtering out less informative features, thereby enabling one to focus on the most informative features for delineating the heterogeneous structure of cells.
\par 
Several methods for selecting HVGs have been proposed. The scran method \cite{lun2016step} uses log-normalized expression values to fit a locally weighted scatterplot smoother (LOESS), which characterizes gene expression variance as a function of the corresponding mean. Each gene's technical variance, represented by the fitted curve, is subtracted from its observed variance to obtain its biological variance. The biological variances are used to determine HVGs.  An alternative option for scran is to use simulated Poisson counts to estimate technical variance. Genes exhibiting observed variance larger than technical variance are designated as HVGs. Seurat version 1 and 2 (Seurat v1, Seurat v2) \cite{satija2015spatial, butler2018integrating} select genes with high dispersion using normalized expression counts, employing different dispersion calculation methods. Cell Ranger \cite{wolf2018scanpy,zheng2017massively} identifies HVGs based on high dispersion calculated from log-normalized expression values. Seurat version 3 (Seurat v3) \cite{stuart2019comprehensive} fits a log-variance against a log-mean curve using raw count values. Additionally, scTransform (SCT) \cite{hafemeister2019normalization} employs a regularized negative binomial regression model for variance stabilizing transformation. The model outputs Pearson residuals. HVGs are selected based on high variance of these residuals.

The HVGs identified by different methods often exhibit significant discrepancies, leading to variations in their effectiveness in revealing the heterogeneity structure of cells (Figure \ref{fig:overview}a). Yet, there is limited knowledge regarding the optimal method in practical applications. A comprehensive understanding of these methods' relative performance necessitates a systematic benchmark study. However, conducting such a study is challenging for multiple reasons. Firstly, it requires a large collection of datasets containing either full or partial ground truth information to ensure robust method evaluation. Secondly, multiple objective evaluation criteria are needed to measure the performance of various methods effectively.
Lastly, the process involves running and comparing a substantial number of methods, adding to the complexity of the study. 

A previous study has compared seven HVG selection methods \cite{yip2019evaluation}, but it did not include a comparison of all commonly used methods. Notably, it omitted the Poisson version of scran and more recent techniques such as Seurat v3 and SCT. Furthermore, the study's scope was limited as it relied on only five benchmark datasets, each containing cells ranging from 92 to 4,296 . The sizes and complexities of these datasets are notably below those of many current scRNA-seq datasets. Another critical limitation of the study is its use of computationally annotated cell type labels as ground truth for method evaluation. 
Since cell type annotation itself involves HVG selection, there exists a risk of bias in the benchmark study, as it may favor the HVG method used to derive the annotations. This ``double dipping'' issue and its resulting bias can limit and potentially undermine the reliability and generalizability of the benchmark results \cite{kriegeskorte2009circular,taylor2015statistical}. 
Consequently, this previous study fell short of comprehensiveness, robustness and generalizability. 


Considering the pivotal role of HVG selection in scRNA-seq analysis, there remains a critical need for a systematic benchmark study of HVG selection methods. To address this gap, we conduct a comprehensive benchmark analysis encompassing 47 HVG selection methods. This includes 21 established and new baseline methods alongside 26 new hybrid methods. Our study utilizes 19 benchmark datasets, each containing full or partial ground truth information, and employs an average of 18 evaluation criteria per method. Our primary contributions can be summarized in three key aspects.

First, we created an extensive benchmark resource consisting of 19 diverse benchmark datasets alongside 18 benchmark metrics. Instead of relying on computational cell type annotations, our benchmark data and metrics incorporate orthogonal information, ensuring an objective, fair, and robust method comparison. This resource can support new method evaluations within the research community.

Second, we categorized HVG selection methods based on their approaches to data transformation and mean-variance adjustment. Employing this framework, we systematically evaluated 7 established methods with publicly available software tools and 14 new methods offering new combinations of data transformation and mean-variance adjustment procedures. This results in the most comprehensive benchmark study of 21 baseline HVG selection methods.

Third, we developed and assessed a novel hybrid strategy that combines multiple baseline methods. By evaluating 26 hybrid methods, we consistently observed superior performance compared to baseline methods, offering an enhanced solution for HVG selection. We have packaged this approach into an R package called \texttt{mixhvg}, which provides both baseline and hybrid methods, with the top-performing hybrid method set as the default for its primary function \texttt{FindVariableFeaturesMix}. Users can seamlessly integrate this package into their analysis pipelines, such as replacing the function \texttt{FindVariableFeatures} in Seurat, for convenient HVG selection.


\begin{figure}[ht]
    \centering
    \includegraphics[width=12.4cm]{Figure1.pdf}
    %%\captionsetup{width=0.96\textwidth}
    \caption{Overview of the benchmark study. (a) An example illustrating the variations in the effectiveness of different HVG selection methods in revealing the heterogeneity structure of cells is shown in the UMAPs of the zheng\_pbmc dataset. The top 2,000 HVGs selected by three methods (random, Seurat v3, and mixHVG (default)) resulted in varying accuracy for clustering cells by cell type, as indicated by the adjusted rand index (ARI). Among these methods, mixHVG performed the best. (b) The benchmark study consists of the following steps: (1) Compiling benchmark data including 7 cell sorting datasets, 7 CITE-seq datasets, and 5 multiomeATAC datasets; (2) Data preprocessing; (3) Compiling 21 baseline methods for selecting HVGs and categorizing them based on data transformation and mean-variance adjustment approaches; (4) Developing 18 objective evaluation criteria (6 for cell sorting data, 6 for CITE-seq data, 6 for multiomeATAC data); (5) Ranking baseline HVG selection methods based on their average performance across all datasets and criteria; (6) Building 26 new hybrid methods using mixtures of baseline methods; (7) Comparing the baseline and hybrid methods for selecting HVGs; (8) Identifying the best methods and making recommendations. }
    \label{fig:overview}
\end{figure}


\section*{Results}
\subsection*{A Computational Resource for Benchmarking HVG Selection Methods}

To facilitate method evaluation, we have constructed a comprehensive benchmark resource consisting of benchmark data, criteria, and a computational pipeline for generating benchmark results (Figure \ref{fig:overview}b). The benchmark pipeline, named benchmarkHVG, has been made openly accessible and is available on GitHub. It can aid developers in evaluating new methods for selecting HVGs in future studies.

Our primary objective is to identify the HVG selection method that most effectively captures cell heterogeneity (i.e., cell-to-cell variation), which constitutes the primary goal of most scRNA-seq experiments. A significant hurdle lies in the scarcity of curated datasets equipped with ground truth information necessary for accurate evaluation. To tackle this challenge, we have compiled 19 datasets, ranging in size from 2,526 to 261,729 cells, for which orthogonal information from other data modalities corresponding to the same cell is available to facilitate method assessment.

The benchmark data comprises 7 datasets from scRNA-seq cell sorting experiments, providing known cell type labels (referred to as ``cell sorting''), 7 CITE-seq datasets offering both cell surface protein and transcriptome information (referred to as ``CITE-seq''), and 5 single-cell multiome datasets that include paired scRNA-seq and scATAC-seq data for each cell (referred to as ``multiomeATAC''). Table~\ref{tab:datatable} summarizes all 19 datasets, including their data types, dataset names, cell counts, gene counts, and protocols. Additionally, the table shows the counts of unique cell type labels for cell sorting datasets, the counts of antibody-derived tags (ADT) representing cell surface proteins for CITE-seq datasets, and the dimension of latent semantic indexing (LSI) used for multiomeATAC datasets.

\newcolumntype{C}[1]{>{\centering\arraybackslash}p{#1}}

\begin{table}[ht]
 
    \caption{List of benchmark datasets.}
    \label{tab:datatable}
    %\begin{adjustbox}{width=\textwidth,center}
    \begin{tabular}{ccccC{0.05\textwidth}C{0.06\textwidth}C{0.03\textwidth}c}
    \hline
        Data Type & Dataset Name & \#Cell & \#Gene & \#Label	& \#ADT & \#LSI & Protocol \\ \hline
        \multirow{8}{*}{Cell Sorting} & GBM\_sd & 3,400 & 23,368 & 3 & & & Smart-seq2 \\ %\hline
        & duo4\_pbmc & 3,994 & 15,568 & 4 & & & 10x Genomics \\ %\hline
        & duo8\_pbmc & 3,994 & 15,716 & 8 & & & 10x Genomics \\ %\hline
        & duo4un\_pbmc	& 6,498 & 16,443 & 4 & & & 10x Genomics \\ %\hline
        & mus\_tissue & 50,816 & 21,458 & 49 & & & Microwell-seq \\ %\hline
        & zheng\_pbmc & 54,388 & 13,435 & 9 & & & 10x Genomics \\ %\hline
        & homo\_tissue	& 131,405 & 18,048 & 39 & & & Microwell-seq \\ %\hline
        %& mus\_tissue\_large & 215,792 & 19,517 & 50 & & & Microwell-seq \\
        \hline
        \multirow{7}{*}{CITE-seq} & pbmc\_cite & 7,665 & 17,014 & & 10 & & \multirow{6}{*}{10x Genomics} \\ %\hline
        & cbmc8k\_cite	& 7,996 & 20,400 & & 13 & & \\ %\hline
        & FBM\_cite & 8,978 & 36,601 & & 198 & & \\ %\hline
        & FLiver\_cite	& 27,912 & 24,247 & & 198 & & \\ %\hline
        & bmcite	& 30,672 & 17,009 & & 25 & & \\ %\hline
        & seurat\_cite	& 161,764 & 20,729 & & 228 & & \\ %\hline
        & sucovid\_cite & 261,729 & 15,114 & & 192 & & \\ \hline
        \multirow{6}{*}{
        $\begin{array}{cc}
            \text{Multiome}\\
            \text{ATAC} 
        \end{array}$} & pbmc3k\_multi & 2,526 & 36,601 & & & 29 & \multirow{6}{*}{10x Genomics} \\ 
        & homo\_brain3k\_multi & 2,896 & 36,601 & & & 29 & \\
        & mus\_brain5k\_multi	& 4,483 & 32,285 & & & 29 & \\ 
        & pbmc10k\_multi & 10,531 & 36,601 & & & 29 & \\ %\hline
        & lymphoma\_multi & 12,861 & 36,601 & & & 29 & \\ \hline
    \end{tabular}
    %\end{adjustbox}
    
    \raggedright{Note: the data types include cell sorting (with discrete cell type labels), CITE-seq (with continuous ADT measurement), multiomeATAC (single-cell multiome data with paired scRNA-seq and scATAC-seq, with scATAC-seq transformed to a low-dimensional cell representation by LSIs).} 
\end{table}

\par 
\subsection*{A Framework for Categorizing HVG Selection Methods}

HVG selection methods can be categorized into four groups based on the format of input data transformation. These groups are determined by whether the methods analyze raw read counts (ct), normalized read counts (nc), log-normalized read counts (lognc), or PFlogPF values, which involve proportional fitting prior to log-transformation followed by additional proportional fitting (PFlogPF) \cite{booeshaghi2022depth}.

Additionally, depending on the criteria for defining ``highly variable'' with respect to the mean-variance relationship adjustment, HVG selection methods can be classified into four categories:
\begin{itemize}
\item \textbf{Mean-variance-based (mv) methods}, which model variance as a function of the mean and select genes with high variance after mean adjustment.
\item \textbf{Log-mean-log-variance-based (logmv) methods}, similar to mv methods, but they model log-variance as a function of log-mean.
\item \textbf{Dispersion-based (disp) methods}, which rank genes based on dispersion, defined as the ratio of variance to mean.
\item \textbf{Mean-max-based (mean\_max) methods}, which choose genes with the highest average expression across cells, as high expression often correlates with high variance. 
%Mean-max-based methods do not consider the relationship between mean and variance.
\end{itemize}

\par

This framework identifies 16 potential classes of HVG selection methods (Table~\ref{tab:method}). Among the established methods, Seurat v1 and v2 belong to dispersion-based HVG selection using normalized counts (disp\_nc\_seuratv1, mvp\_nc\_seuratv2). Seurat v3 is a log-mean-log-var-based  approach based on raw counts (logmv\_ct\_seuratv3). Scran provides a mean-variance-based method based on log-normalized counts (mv\_lognc\_scran).
Scanpy \cite{wolf2018scanpy} offers three options for HVG selections, including Seurat v1, v3, and a cell-ranger-based one. We applied Seurat v1 and v3 in the original implementation, but used cell-ranger HVG selection with scanpy, which adopts a dispersion-based method using log-normalized counts (scanpy\_cell\_ranger). Since the existing methods do not cover all scenarios, we developed 13 additional methods to cover all 16 classes (Table~\ref{tab:method}). Specifically, we adapted the frameworks of scran, Seurat v3, and Seurat v1 to implement mean-variance-based, log-mean-log-var-based, and dispersion-based methods, for different types of input data transformation, respectively.

SCT stands as a method outside this classification. It uses an integrated approach where HVG selection and data transformation are inseparable and ranks genes based on Pearson residuals following variance stabilization. Scran also offers an alternative method that estimates technical variance using simulated Poisson counts and subtracts this from the observed variance to describe genes' biological variance. Furthermore, we introduced a negative control approach that randomly selects nonzero-expressed genes, i.e., genes with nonzero expression in at least one cell. These components lead to a total of 21 baseline methods in our benchmark study.

\begin{table}[ht]
\caption{Classification of baseline HVG selection methods.}
    \label{tab:method}
\begin{tabular}{@{\extracolsep\fill}C{0.15\textwidth}C{0.15\textwidth}C{0.17\textwidth}C{0.2\textwidth}C{0.16\textwidth}}
\toprule%
\multicolumn{5}{l}{Methods classified based on input data transformation and mean-variance adjustment approaches} \\
\midrule
& mean-var & $\begin{array}{c}\text { log-mean- } \\
\text { log-var }\end{array}$ & dispersion & mean\_max \\
\midrule
Count & \text{mv\_ct} & \textbf{logmv\_ct\_seuratv3} & disp\_ct & mean\_max\_ct \\
& & & & \\
$\begin{array}{c}\text { Normalized } \\
\text { Count }\end{array}$ & mv\_nc & logmv\_nc & $\begin{array}{c}\textbf{disp\_nc\_seuratv1} \\
\textbf{mvp\_nc\_seuratv2}\end{array}$ & mean\_max\_nc \\
& & & & \\
%\hdashline
$\begin{array}{c}\text { LogNormalized } \\
\text { Count }\end{array}$ & \textbf{mv\_lognc\_scran} & logmv\_lognc & $\begin{array}{c}\textbf{scanpy\_cell\_ranger}\\ \text{disp\_lognc}\end{array}$ & mean\_max\_lognc \\
& & & & \\
PFlogPF & mv\_PFlogPF & logmv\_PFlogPF & disp\_PFlogPF & mean\_max\_PFlogPF \\
\midrule
\multicolumn{5}{l}{Other non-classified methods} \\
\multicolumn{5}{c}{\textbf{SCT},  \textbf{poisson\_scran}, \text{random}} \\
\botrule
\end{tabular}
\footnotetext{Note: existing methods with software tools are shown in bold.}
\end{table}


\subsection*{Benchmark Baseline Methods Using Known Cell Type Labels}

Our evaluation of the 21 baseline methods begins with the 7 cell sorting datasets (Table~\ref{tab:datatable}).  These datasets contain mixtures of sorted cells, where the true cell type labels are obtained from cell sorting experiments. For instance, Figures~\ref{fig:sort}a and \ref{fig:sort}b illustrate the zheng\_pbmc dataset, which consists of sorted cells from a peripheral blood mononuclear cell (PBMC) sample. Figure~\ref{fig:sort}a displays cells grouped by Louvain clustering and color-coded according to their cluster labels, while Figure~\ref{fig:sort}b colors cells by their true cell type labels.


\begin{figure}[htbp]
    \centering
    \includegraphics[width=\textwidth]{Figure2.pdf}
    %\captionsetup{width=0.96\textwidth}
    \caption{Evaluation of baseline methods based on cell sorting datasets. \textbf{(a)} Cells in the zheng\_pbmc dataset shown in UMAP. Different colors indicate cell clusters identified by Louvain clustering, performed using the top 30 principal components of HVGs selected by Seurat v3. \textbf{(b)} The same UMAP as in (a), but with colors indicating true cell type labels. \textbf{(c)}-\textbf{(g)} schematic diagrams of six evaluation criteria for benchmark datasets with discrete true cell type labels: \textbf{(c)} Adjusted Rand Index (ARI) and Normalized Mutual Information (NMI); \textbf{(d)} The ratio of between-cell-type variance to within-cell-type variance; \textbf{(e)} $k$-NN classification accuracy based on true labels; \textbf{(f)} Average Silhouette Width (ASW); \textbf{(g)} Cell-type Local Inverse Simpson's Index (LISI). \textbf{(h)} Performance of 21 baseline HVG selection methods across six evaluation criteria. Each row represents a method, and each column represents a criterion. For each method and criterion, the average performance across 7 cell sorting datasets is shown in a balloon plot. Methods are ranked based on each criterion. The bar plot on the left displays the average rank across the 6 criteria. Existing HVG methods with publicly available software tools are marked with an asterisk (*). Those without a star are new methods we implemented to make the benchmark study comprehensive.}
    \label{fig:sort}
\end{figure}

We reasoned that improved HVG selection should better characterize cell heterogeneity, resulting in improved cell clustering and better separation of distinct cell types. 
Accordingly, we evaluated the methods using six criteria: 
\begin{itemize} 
\item \textbf{Adjusted Rand Index (ARI)} and 
\item \textbf{Normalized Mutual Information (NMI)}: Both measuring the consistency between HVG-based cell clustering and true cell type labels (Figure~\ref{fig:sort}c).
\item \textbf{Variance Ratio}: The ratio of between-cell-type variance to within-cell-type variance, measuring the separation of different cell types (Figure~\ref{fig:sort}d). 
\item \textbf{$k$-nearest-neighbor ($k$-NN) classification accuracy}: The accuracy for predicting a cell’s type based on the cell type labels of its $k$-nearest neighbor cells, reflecting the clustering of similar cells (Figure~\ref{fig:sort}e).
\item \textbf{Average Silhouette Width (ASW)}: Measuring the separation of different cell types by comparing the average distance of a cell to others of the same cell type versus the average distance to cells from the closest other cell type (Figure~\ref{fig:sort}f).
\item \textbf{Cell-type Local Inverse Simpson Index (LISI)}: Characterizing the local diversity of cell types around a cell, calculated by the expected number of draws needed to obtain at least one cell with the same cell type from a cell's neighbors (Figure~\ref{fig:sort}g).
\end{itemize}
For consistent interpretation, all six criteria were transformed (e.g., LISI was inverted to 1/LISI) so that larger values indicate better performance. For a fair method comparison, the top 2,000 HVGs from each method were used for downstream analyses, consistent with Seurat’s default configuration. To demonstrate, Additional file 1: Supplementary Figure~S1 compares the seven existing HVG methods in the zheng\_pbmc dataset using the six criteria along with cells' UMAP plots. In this dataset, Seurat v3 (logmv\_ct\_seuratv3) did not separate cytotoxic and naive cytotoxic cells well (see red circle). Consistent with this observation, the performance metrics of Seurat v3 did not have the largest values. Similarly, Seurat v2 (mvp\_nc\_seuratv2), Cell ranger (scanpy\_cell\_ranger) and SCT also did not perform well. By contrast, cytotoxic and naive cytotoxic cells showed better separation in scran (mv\_lognc\_scran, poisson\_scran) and Seurat v1 (disp\_nc\_seuratv1). Correspondingly, the performance metrics of these methods showed higher values.

After ranking methods for each criterion and each dataset, we compared the average rank of each method across all datasets and criteria (Figure~\ref{fig:sort}h). 
All HVG methods performed better than randomly selected genes, highlighting the effectiveness of HVG selection strategies. Overall, the Poisson-based scran (poisson\_scran) showed the best performance, followed by the mean-max-based methods using normalized counts (mean\_max\_nc) and raw counts (mean\_max\_ct)  (Figure~\ref{fig:sort}h).  
In this evaluation, mean-max-based methods generally outperformed other approaches like dispersion, mean-variance, and log-mean-log-var methods. Within each method for defining ``highly variable" (i.e., mean-max, dispersion, log-mean-log-var, or mean-var), the optimal input data transformation varied. For instance, for mean-max-based methods, normalized count (nc) performed the best. For dispersion-based methods, raw count (ct) was optimal. For log-mean-log-var-based methods, log-normalized count (lognc) was the best. For mean-var-based methods, PFlogPF outperformed the others. 


%We illustrate the evaluation criteria with schematic diagrams in Figure~\ref{fig:sort}(c,d,e,f) for discrete labels, and Figure~\ref{fig:cite}(d,e,f,g) for continuous information. All the displayed criteria/ranks in Figure~\ref{fig:sort}(g), \ref{fig:cite}(h), \ref{fig:multi}(d), \ref{fig:cmp}(a) follow increasing trend: higher/larger is better. In cases where original criteria/ranks follow the opposite direction, we invert them to maintain consistency. For example, knn\_mse($k$-nearest neighbors mean square error) achieves the best performance when minimized, so we use inv\_knn\_mse. Bar plots in Figure~\ref{fig:sort}(g), \ref{fig:cite}(h), \ref{fig:multi}(d), \ref{fig:cmp}(a) rank the 21 baseline methods correspondingly. Initially, we rank the 18 methods for each criterion and each dataset, and then average over all criteria and datasets. Finally, we display the inverse of average ranks to represent better performance as an increasing trend. Dot plots in Figure~\ref{fig:sort}(g), \ref{fig:cite}(h), \ref{fig:multi}(d), \ref{fig:cmp}(a) average over all datasets for each criterion: larger circles/lighter colors are better. In particular, in Figure~\ref{fig:cmp}(a), color is scaled for each criterion, while circle size is scaled across all criteria. We annotate three top-performing methods for each criterion and append a star (*) after the names of widely used publicly available HVG methods.


\subsection*{Benchmark Baseline Methods Using CITE-seq}

We then proceeded to evaluate the 21 baseline methods across 7 CITE-seq datasets (Table~\ref{tab:datatable}). Each dataset includes both scRNA-seq data and antibody-derived tag (ADT) measurements for surface proteins. The number of ADTs ranges from 10 to 228. Figures~\ref{fig:cite}a and \ref{fig:cite}b use the FLiver\_cite dataset as an example. In Figure~\ref{fig:cite}a, cells are color-coded by clusters obtained using scRNA-seq and Louvain clustering. Figure~\ref{fig:cite}b displays cells color-coded based on clusters obtained using surface protein (ADT) profiles.


\begin{figure}[htbp]
    \centering
    \includegraphics[width=0.88\textwidth]{Figure3.pdf}
    %\captionsetup{width=0.96\textwidth}
    \caption{Evaluation of baseline methods based on CITE-seq. \textbf{(a)}-\textbf{(b)} Cells in the cbmc8k\_cite dataset shown in UMAP generated using scRNA-seq data. 
    Colors indicate cell clusters identified by (a) scRNA-seq using top 30 principal components of HVGs selected by Seurat v3, and (b) surface protein (ADT) profiles.
    \textbf{(c)} Scatter plot comparing scRNA-seq HVG-based and ADT-based cell-to-cell distances. \textbf{(d)}-\textbf{(h)} Schematic diagrams of six evaluation criteria for benchmark datasets with orthogonal data modalities taking continuous values: \textbf{(d)} The ratio of between-cluster-variance to within-cluster-variance of ADT values; \textbf{(e)}  $k$-NN regression mean square error (MSE) for predicting ADT; \textbf{(f)} The ratio of non-$k$-nearest-neighbor distance to $k$-nearest-neighbor distance; \textbf{(g)} Cell-to-cell distance correlation between scRNA-seq and ADT; \textbf{(h)} Adjusted Rand Index (ARI) and Normalized Mutual Information (NMI). \textbf{(i)} Performance of 21 baseline HVG methods across six evaluation criteria. The balloon plot shows average performance across 7 CITE-seq datasets. Methods are ranked based on each criterion. The bar plot on the left displays the average rank across the 6 criteria.}
    \label{fig:cite}
\end{figure}

We reasoned that improved HVG selection should more effectively characterize cell heterogeneity and cell-to-cell similarity. Enhanced cell-to-cell similarity measurements derived from RNA-seq data should align more closely with cells’ similarity based on surface protein (ADT) profiles (Figure~\ref{fig:cite}c).
Here, protein (ADT) profiles offer orthogonal information for evaluating scRNA-seq analyses. Unlike the cell sorting data with discrete cell type labels, the ADT profiles in CITE-seq data take continuous values. Accordingly, we evaluated methods in six different ways: 
\begin{itemize}
\item \textbf{Variance ratio}: Cell clustering is based on scRNA-seq gene expression profiles of HVGs, and the ratio of between-cluster to within-cluster variance using ADT profiles is calculated to assess the separation of cells’ ADT profiles (Figure~\ref{fig:cite}d).
\item \textbf{$k$-NN regression mean squared error (MSE)}: The accuracy of predicting a cell’s ADT profile using its $k$-nearest neighbors, identified by the gene expression of HVGs (Figure~\ref{fig:cite}e).
\item \textbf{$k$-NN distance ratio}: Measuring the average ADT distance to $k$-nearest neighbors against the average ADT distance to all other cells, with neighbors determined by HVG expression in scRNA-seq (Figure~\ref{fig:cite}f).
\item \textbf{Distance correlation}: Assessing the Pearson correlation between the ADT-based cell-to-cell distance and the HVG expression-based cell-to-cell distance (Figure~\ref{fig:cite}g).
\item \textbf{Adjusted Rand Index (ARI)} and
\item \textbf{Normalized Mutual Information (NMI)}: Both quantifying the consistency between cell clustering according to HVG expression and cell clustering by ADTs (Figure~\ref{fig:cite}h).
\end{itemize}
All six criteria were appropriately adjusted (e.g., transforming MSE to 1/MSE) so that higher values indicate superior performance. To demonstrate, Additional file 1: Supplementary Figure~S2 compares the seven existing HVG methods in the cbmc8k\_cite dataset. Based on the six criteria, scran (mv\_lognc\_scran, poisson\_scran) and Seurat v1 (disp\_nc\_seuratv1) outperformed Seurat v3 (logmv\_ct\_ seuratv3), Seurat v2 (mvp\_nc\_seuratv2), Cell ranger (scanpy\_cell\_ranger) and SCT in this dataset. Indeed, the ADT cluster 1 and cluster 8 were not separated well in scRNA-seq space using HVGs selected by Seurat v3, Seurat v2, Cell ranger and SCT, but they formed distinct clusters using HVGs selected by scran and Seurat v1.

\par 
We ranked all HVG selection methods across the six criteria and the CITE-seq datasets, as shown in Figure~\ref{fig:cite}i. Once again, every method we tested was more effective than choosing genes at random. The dispersion and normalized count-based method Seurat v1 (disp\_nc\_seuratv1) performed the best, followed by two mean-var-based methods on normalized count (mv\_nc) and log-normalized count (mv\_lognc\_scran). In this evaluation, mean-var-based methods showed better overall performance than mean-max, log-mean-log-var, and dispersion-based methods, except for disp\_nc\_seuratv1, which was the best. Within each method for defining ``high variability", such as mean-var, log-mean-log-var, dispersion, or mean-max, the best results were achieved using either normalized counts (nc) or log-normalized counts (lognc) as input data types. In contrast, other data types like PFlogPF and raw counts did not perform as well in our evaluation.


\subsection*{Benchmark Baseline Methods Using Paired scRNA-seq and scATAC-seq}

We further extended our analysis using 5 single-cell multiome datasets, which provide paired scRNA-seq and scATAC-seq data (multiomeATAC), to evaluate the 21 baseline methods (Table \ref{tab:datatable}). Similar to CITE-seq, scATAC-seq provides orthogonal information for evaluating cell heterogeneity characterized by scRNA-seq. Latent semantic indexing (LSI) \cite{deerwester1990indexing} was used to transform scATAC-seq data into a low-dimensional embedding format. 

Figures~\ref{fig:multi}a and \ref{fig:multi}b show the pbmc3k\_multi dataset as an example, where cells are color-coded based on their clustering obtained from scRNA-seq using principal components of HVGs and scATAC-seq using LSIs, respectively. For scATAC-seq, we disregarded the first LSI dimension because it predominantly reflects sequencing depth, which is a technical variance rather than a biological difference (Additional file 1: Supplementary Figure~S3). Instead, we used the $2^{\text{nd}}$ to $30^{\text{th}}$ dimensional LSIs to calculate similarities between cells based on scATAC-seq data. Our findings show a correlation between cell-to-cell similarities computed from scATAC-seq and scRNA-seq data, as illustrated in Figure~\ref{fig:multi}c. 

We reasoned that improved HVG selection should refine scRNA-seq-derived cell-to-cell similarity which would in turn better align with scATAC-seq-based cell-to-cell similarity.
For the multiome data evaluation, we applied the same six criteria used for CITE-seq analysis, with the only difference being that LSIs were not scaled for distance calculation.
To demonstrate, Additional file 1: Supplementary Figure~S4 compares the seven existing HVG methods in the pbmc10k\_multi dataset. Here, Seurat v3 showed the best overall performance based on the six evaluation criteria.

\par 
Based on each method's average rank across multiomeATAC datasets and evaluation criteria (Figure~\ref{fig:multi}d), all methods outperformed randomly selected genes. The log-mean-log-var-based method Seurat v3 (logmv\_ct\_seuratv3) performed the best, followed by two mean-var-based methods on log-normalized count (mv\_lognc\_scran)  and normalized count (mv\_nc). Overall, mean-var-based methods showed better performance than mean-max, dispersion, and log-mean-log-var-based methods, except for logmv\_ct\_seuratv3, which was the best. Within each way for defining ``high variability'' (mean-max, dispersion, log-mean-log-var, or mean-var), the optimal input data type varied. For mean-max and dispersion-based methods, normalized count (nc) was optimal. For log-mean-log-var and mean-var-based methods, raw count (ct) and log-normalized count (lognc) were optimal, respectively. 


\begin{figure}[htbp]
    \centering
    \includegraphics[width=\textwidth]{Figure4.pdf}
    %\captionsetup{width=0.96\textwidth}
    \caption{Evaluation of baseline methods based on single-cell multiome datasets with paired scRNA-seq and scATAC-seq. \textbf{(a)}-\textbf{(b)} Cells in the pbmc10k\_multi dataset shown in UMAP generated using scRNA-seq data. 
    Colors indicate cell clusters identified by (a) scRNA-seq and Louvain clustering using the top 30 principal components of HVGs selected by Seurat v3, and (b) the $2^{\text{nd}}$ to $30^{\text{th}}$ LSIs of scATAC-seq.
    \textbf{(c)} Scatter plot comparing scRNA-seq HVG-based cell-to-cell distances and scATAC-seq-based cell-to-cell distance. HVGs are selected by Seurat v3.
    \textbf{(d)} Performance of 21 baseline HVG selection methods across six evaluation criteria. The average performance across 5 single-cell multiome datasets is shown in a balloon plot. Methods are ranked based on each criterion. The bar plot on the left displays the average rank across the 6 criteria.}   
    \label{fig:multi}
\end{figure}


\subsection*{Joint Evaluation of Baseline Methods}
Combining the results from cell sorting, CITE-seq and single-cell multiomeATAC datasets, an overall rank was assigned to each method (Figure~\ref{fig:cmp}a), leading to the following conclusions.

\begin{figure}[htbp]
    \centering
    \includegraphics[width=0.95\textwidth]{Figure5.pdf}
    %\captionsetup{width=0.96\textwidth}
    \caption{Comparison of 21 baseline HVG selection methods across all benchmark datasets and criteria. 
    \textbf{(a)} Overall ranking of the methods. The methods are categorized into four main groups based on the mean-variance-adjustment approach: mean-var-based, log-mean-log-var-based, dispersion-based, and mean-max-based. Each category is represented by a different color. Methods that do not fit into these categories are colored separately. Existing methods with publicly available software tools are indicated with an asterisk (*). The evaluation involves 18 criteria grouped based on benchmark data types: cell sorting, CITE-seq, and multiomeATAC. Each data type has 6 evaluation criteria. For each method and criterion, the average performance across all benchmark datasets is shown in a balloon plot. Methods are ranked based on each criterion. 
    The bar plot on the left displays the average rank across the 18 criteria. Taller bars, bigger balloons, or lighter balloon colors indicate better performance. The top-performing methods include poisson\_scran, mv\_lognc\_scran, mv\_nc and disp\_nc\_seuratv1.
    \textbf{(b)} A heatmap displaying the overlap among the top 2,000 HVGs selected by the 21 baseline methods in the pbmc3k\_multi dataset. Different mean-variance adjustment method categories showed little overlap in their selected HVGs.}
    \label{fig:cmp}
\end{figure}

First, our analysis confirmed that all HVG selection methods outperformed randomly selected genes, underscoring the importance of HVG selection for accurately studying cell heterogeneity.

Second, we observed that the most effective method varied depending on the type of benchmark data and specific criteria used. No single HVG selection method consistently outperformed the others across all criteria and datasets. For instance, while the Poisson-based method poisson\_scran performed the best in cell sorting benchmark (Figure~\ref{fig:sort}h), the dispersion and normalized count-based method, Seurat v1 (disp\_nc\_seuratv1), excelled in the CITE-seq benchmark (Figure~\ref{fig:cite}i), and the log-mean-log-var-based method, Seurat v3 (logmv\_ct\_seuratv3), ranked the top in single-cell multiomeATAC benchmark (Figure~\ref{fig:multi}d). Within the single-cell multiomeATAC data, while the mean-var-based method on count (mv\_ct) performed the best in terms of variance ratio, the mean-var-based method on log-normalized count (mv\_lognc\_scran) excelled in terms of the $k$-NN distance ratio and $k$-NN regression MSE, and log-mean-log-var-based method (logmv\_ct\_seuratv3) was the top method in terms of distance correlation and ARI (Figure~\ref{fig:multi}d).

Third, when considering the average ranks across all datasets and criteria, the four top-performing methods were: (1) poisson\_scran: scran based on Poisson modeling of technical variance; (2) mv\_lognc\_scran: scran based on modeling mean-variance curves of log-normalized counts; (3) mv\_nc: modeling mean-variance curves of normalized counts; and (4) disp\_nc\_seuratv1: Seurat v1 based on dispersion of normalized counts (Figure~\ref{fig:cmp}a). 
These methods demonstrated competitive overall performance across various benchmark data types and criteria. In contrast, some approaches, like mean-max-based methods, performed well in one data type, e.g., cell sorting, but did not perform as well in others, indicating that they may not be universally competitive.


Finally, upon comparing methods in different categories (as color-coded in Figure~\ref{fig:cmp}a), mean-max-based and log-mean-log-var-based methods were generally less effective compared to mean-var-based methods. 

We also compared method categories from the view of data transformation (Additional file 1: Supplementary Figure~S5). No data transformation uniformly outperformed the others, except that normalized counts (nc) consistently performed better than PFlogPF within each mean-variance adjustment method category (Figure~\ref{fig:cmp}a).


\par 

\subsection*{Hybrid Methods Significantly Enhance HVG Selection}
Comparisons of HVGs reveal that different methods identify distinct gene sets (Figure~\ref{fig:cmp}b, Additional file 1: Supplementary Figure~S6), indicating that various methods may capture different aspects of the dataset. Pairwise comparisons within and between different method categories further demonstrate that methods within the same mean-variance adjustment category tend to select more similar HVGs than those from different categories. For instance, HVGs selected by different mean-max-based methods exhibit considerable overlap, although with minimal overlap with HVGs identified by dispersion-based methods.


\begin{figure}[htbp]
    \centering
    \includegraphics[width=\textwidth]{Figure6.pdf}
    %\captionsetup{width=0.96\textwidth}
    \caption{Comparison of baseline methods and hybrid methods for selecting HVGs. \textbf{(a)} Illustration of hybrid approach. In this toy example, baseline methods 1 and 2 each generate a ranked list of genes. The hybrid approach assigns a score to each gene based on its best rank from methods 1 and 2, and then reranks all genes accordingly. Here, the hybrid method identifies gene A and gene D as the top two HVGs.
    \textbf{b} Evaluation of 26 hybrid HVG selection methods and 21 baseline methods across all benchmark data and criteria. For all 47 methods, the top 2,000 HVGs are used. The average performance across all benchmark datasets is depicted in a balloon plot for each method and evaluation criterion. Methods are ranked based on each criterion. The bar plot on the left illustrates the average rank across all 18 criteria. Hybrid methods are color-coded according to the number of baseline methods they incorporate, with each hybrid method named based on the mixture of baseline methods used. The three top-performing methods and the five baseline methods used to construct hybrid methods are highlighted by red dashed lines in the balloon plot. 
    }
    \label{fig:cmp2}
\end{figure}

Prompted by these observations, we explored whether combining two or more HVG selection methods could more effectively capture dataset information than individual methods. To investigate this, we developed and evaluated 26 hybrid methods, each combining two or more baseline HVG selection methods. We considered various combinations of five baseline methods:  mv\_lognc\_scran (1mv), logmv\_lognc (2lmv), poisson\_scran (3pos), disp\_nc\_seuratv1 (4dis), and mean\_max\_nc (5max). These included the top-performing baseline methods from each mean-variance-adjustment method categories: mean-var-based (1mv), log-mean-log-var-based (2lmv), dispersion-based (4dis), and mean-max-based (5max). Additionally, the top-performing method from Figure~\ref{fig:cmp}, poisson\_scran (3pos), was included. 
Notably, mv\_lognc\_scran (1mv), poisson\_scran (3pos), disp\_nc\_seuratv1 (4dis), and the excluded mv\_nc are top four methods in baseline method evaluation (Figure~\ref{fig:cmp}a). We excluded mv\_nc because it yielded similar HVGs as mv\_lognc\_scran (1mv) (Figure~\ref{fig:cmp}b). 

To comprehensively evaluate method combinations, we explored all possible mixtures of two, three, four, and five methods, resulting in 26 distinct hybrid method variants. For each hybrid method, genes were ranked using each baseline method in the mixture. A gene's final score was then determined based on its highest rank across these baseline methods, which was used to rerank all genes. Figure~\ref{fig:cmp2}a illustrates our gene selection strategy. Here genes are reranked based on their best and second-best rankings derived from either method 1 or method 2, and genes A and D are identified as the top two HVGs. 


Overall, hybrid methods demonstrated superior performance compared to baseline methods (Figure~\ref{fig:cmp2}b). 
When evaluating all methods together, the 26 hybrid methods achieved significantly better rankings compared to the 20 baseline methods excluding the ``random'' method which is a negative control (one-sided t-test $p\text{-value} < 1\times 10^{-8}$). The improvement remained significant when the hybrid methods were compared to the 5 baseline methods used to construct them (one-sided t-test $p\text{-value}= 0.016$).
Notably, the top three methods in the overall ranking of all 47 methods were all hybrid methods, including (1) a mixture of mv\_lognc\_scran, poisson\_scran and disp\_nc\_seuratv1 (1mv3pos4dis), (2) a mixture of mv\_lognc\_scran, logmv\_lognc and poisson\_scran (1mv2lmv3pos), and (3) a mixture of mv\_lognc\_scran and poisson\_scran (1mv3pos). 

By comparing different methods in individual datasets, we observed that while some baseline methods showed optimal performance in some datasets, no baseline method consistently performed among the top across all datasets. In contrast, the optimal hybrid approach robustly showed competitive performance, performing either among the top or close to the top, across different datasets, explaining its superior overall performance (Additional files 1: Supplementary Figures~S1, S2, S4).
%, \ref{fig:cd34}).

Our findings underscore the effectiveness of the hybrid approach in improving HVG selection. To facilitate the utilization of the hybrid approach, we have integrated all 26 HVG method mixtures into the R package, \texttt{mixhvg}.
Users can utilize these methods through calling the \texttt{FindVariableFeaturesMix} function, with 1mv3pos4dis being designated as the default method.

 

\subsection*{Computational Efficiency and Scalability}

The computational efficiency and scalability of various methods were evaluated by assessing CPU time and memory usage on a Dell PowerEdge R6525 server, equipped with 2 AMD EPYC 7713 64-Core CPUs (2.0GHz, 256MB L3 Cache), and 1TB of RAM.
The comparison, including seven existing baseline methods and the top-performing hybrid method (1mv3pos4dis), is illustrated in Figure~\ref{fig:scalability}. Both running time and memory usage demonstrated linear scaling with increasing cell numbers across all methods.

SCT exhibited higher memory requirements and longer running times compared to other methods. Regarding the hybrid method (1mv3pos4dis), if each contributing method is executed sequentially, the computation time approximately equals the sum of the contributing methods' times, and the memory usage is constrained by the largest memory requirement among the individual contributing methods. 
Given the favorable scalability of each individual method, the hybrid approach also exhibits robust scalability and can effectively handle large-scale datasets.
For instance, analyzing a dataset with over 160,000 cells required approximately 120 seconds and 10GB of memory.

\begin{figure}[htbp]
    \centering
    \includegraphics[width=\textwidth]{Figure7.pdf}
    %%\captionsetup{width=0.96\textwidth}
    \caption{Computational efficiency and scalability of HVG selection methods. \textbf{(a)} CPU time and \textbf{(b)} Memory usage for different data sizes (i.e., number of cells). The plot includes seven existing methods with publicly available software tools denoted by an asterisk (*) and the optimal hybrid method 1mv3pos4dis. The datasets utilized for this analysis include pbmc3k\_multi, mus\_brain5k\_multi, cbmc8k\_cite, FLiver\_cite, mus\_tissue, seurat\_cite. They are arranged based on increasing cell numbers.}
    \label{fig:scalability}
\end{figure}

\section*{Discussion and Conclusions}
In summary, we have created a benchmark resource comprising 19 datasets and 6 evaluation criteria per dataset to assess methods for selecting HVGs from scRNA-seq data. Through this tool, we systematically evaluated 47 HVG selection methods, including 21 baseline and 26 hybrid methods, performing 5,358 analyses (47 methods $\times$ 19 datasets $\times$ 6 criteria per dataset). This represents the most comprehensive benchmark study of HVG selection methods to date.

Our study leverages experimental data orthogonal to RNA-seq, such as true cell type labels from cell sorting, protein levels from CITE-seq, and ATAC-seq open chromatin profiles from single-cell multiome (multiomeATAC) data, for objective evaluation of selected HVGs. This approach addresses previous issues of ``double dipping'', where cell type labels used for evaluation are derived from specific HVG methods. This rigorous evaluation framework, diverse benchmark datasets, and multiple evaluation criteria together make our conclusions robust.

Our results demonstrate that both baseline and hybrid HVG selection methods outperform randomly selected genes, highlighting the importance of HVG selection. However, optimal baseline methods vary depending on dataset type and evaluation criteria. Notably, hybrid methods combining multiple baseline methods, robustly outperform individual baseline methods. Considering the significance of HVG selection in scRNA-seq analysis, we advocate for adopting the hybrid approach as the new standard. To facilitate this transition and streamline integration into existing workflows, we developed the R package \texttt{mixhvg}, incorporating evaluated hybrid methods.

Our study's main contributions include a comprehensive evaluation of HVG selection methods, introduction of the hybrid approach to enhance HVG selection, development of \texttt{mixhvg} software to support hybrid HVG selection methods, and creation of a benchmark resource. These resources can aid future evaluations of new HVG methods.

While our focus has been on scRNA-seq cell heterogeneity, the proliferation of single-cell and spatial technologies introduces other data modalities, such as scATAC-seq and spatial transcriptomic and epigenomic data, each with unique characteristics. Future investigations should explore optimal feature selection strategies for these modalities and assess whether hybrid approaches can similarly enhance existing analysis tools and pipelines.


\section*{Methods} 

\subsection*{HVG Selection Methods}
\subsubsection*{Seurat v1 and v2}
Seurat v1 \cite{satija2015spatial} identifies genes with the highest dispersion in normalized read counts. 
Seurat v2 \cite{butler2018integrating} categorizes genes into bins according to their mean expression levels of normalized counts. Within each bin, the dispersion is standardized by calculating a z-score. HVGs are then determined based on the highest standardized dispersion.

\subsubsection*{Seurat v3}
Seurat v3 \cite{stuart2019comprehensive} employs a locally weighted scatterplot smoother (LOESS) to model the relationship between log-transformed variance and log-transformed mean of genes, computed from their raw read counts. This allows for the creation of a standardized expression profile for genes, adjusting for the observed mean and expected standard deviation derived from the LOESS curve. The standardized values are truncated to a maximum value to reduce the impacts of outliers. Genes are then ranked based on the variance calculated from the truncated standardized values.


\subsubsection*{SCT (SCTransform)}
SCT \cite{hafemeister2019normalization} uses a regularized negative binomial regression model for normalizing gene expression data. The observed unique molecular identifier (UMI) counts are converted to Pearson residuals based on the model-estimated means and standard deviations. Genes are then ranked based on the variance of these Pearson residuals.

\subsubsection*{Scran}
Scran \cite{lun2016step} assumes that a gene's total variance comprises both biological and technical components. It fits a variance-mean curve using the log-normalized expression values of genes. For each gene, the curve's fitted value represents an estimate of its technical variance,  which is subtracted from the total variance to isolate the biological component. This biological variance is then utilized to rank genes.
\par 
Additionally, scran offers an alternative method assuming Poisson noise  \cite{lun2016step}. This method first simulates Poisson counts for fake genes across all cells where genes' mean expression and cells' library size are representative of the observed mean expression and library size. It then uses the simulated data to estimate the technical variance, which is subtracted from the observed total variance of each gene to obtain its biological variance.


\subsubsection*{Cell Ranger}
In Cell Ranger \cite{wolf2018scanpy,zheng2017massively}, genes are divided into several bins according to the quantiles of their means. Within each bin, normalized dispersion is computed as the absolute difference between dispersion and the median dispersion, divided by the median absolute deviation of dispersion. Genes exhibiting high normalized dispersion are subsequently selected.

\subsubsection*{Other Baseline HVG Selection Methods}
In Table~\ref{tab:method}, methods available in existing software tools are shown in bold. The other methods were implemented by tailoring existing tools to accommodate different data transformations. For example, mv\_ct, mv\_nc, and mv\_PFlogPF were implemented by using raw counts, normalized counts, and PFlogPF to replace log-normalized counts as input data for the mean-variance-based method mv\_lognc\_scran. Similarly, logmv\_nc, logmv\_lognc, and logmv\_PFlogPF were implemented by replacing the input data type for the log-mean-log-variance-based method Seurat v3. The dispersion-based methods disp\_ct, disp\_lognc, and disp\_PFlogPF were implemented by replacing the input data type for Seurat v1. For the mean\_max family, the mean gene expression across cells computed using the corresponding data transformation was used to rank genes.

\subsubsection*{Hybrid HVG Selection Methods}
Each hybrid method is constructed by aggregating two or more baseline methods as follows. First, genes are ranked by each baseline method. For each gene, its highest (i.e., best) ranking across the baseline methods is assigned as its score. Next, genes are ranked based on this score. If two genes have identical best rankings from the baseline methods, the second-highest rankings are used to break the tie. This process continues with subsequent rankings as needed until the tie is resolved. 

\subsubsection*{Baseline Method Nomenclature}
For brevity, we have adopted the following abbreviations: ct for raw counts, nc for normalized counts, lognc for log-normalized counts, PFlogPF for normalized values derived from proportional fitting prior to log-transformation followed by additional proportional fitting, mv for mean-variance, logmv for log-mean-log-variance, disp for dispersion, and mean\_max for maximum of mean expression.

The 21 baseline HVG selection methods are labelled as follows:
(1) random, (2) mv\_ct, (3) mv\_nc, (4) mv\_lognc\_scran,  (5) mv\_PFlogPF, (6) logmv\_ct\_seuratv3, (7) logmv\_nc, (8) logmv\_lognc, (9) logmv\_PFlogPF, (10) disp\_ct, (11) disp\_nc\_seuratv1, (12) mvp\_nc\_seuratv2, (13) scanpy\_cell\_ranger, (14) disp\_lognc, (15) disp\_PFlogPF, (16) mean\_max\_ct, (17) mean\_max\_nc, (18) mean\_max\_lognc, (19) mean\_max\_PFlogPF, (20) SCT, (21) poisson\_scran. 
Among these methods, (2)-(5) are mean-variance-based (mv), (6)-(9) are log-mean-log-var-based (logmv), (10)-(15) are dispersion-based (disp), (16)-(19) are mean-max-based, (20) is Pearson-residual-based, and (21) is Poisson-based.
Seurat v1 (disp\_nc\_seuratv1), Seurat v2 (mvp\_nc\_seuratv2), Seurat v3 (logmv\_ct\_seuratv3), scran (mv\_lognc\_scran), scran with poisson (poisson\_scran), and Cell Ranger (scanpy\_cell\_ranger) are the seven established methods with publicly available software. 


\subsubsection*{Hybrid Method Nomenclature}
The hybrid methods are derived from combinations of five baseline methods, which are mv\_lognc\_scran (1mv), logmv\_lognc (2lmv), poisson\_scran (3pos), disp\_nc\_seuratv1 (4dis), and mean\_max\_nc (5max). We refer to these methods by their abbreviations: 1mv, 2lmv, 3pos, 4dis, and 5max. When creating hybrid HVG selection methods from any number of these baseline methods, we name the hybrid method using a concatenation of the selected methods' abbreviations. For example, a hybrid method combining mv\_lognc\_scran, poisson\_scran, and disp\_nc\_seuratv1 is named 1mv3pos4dis.
\par 


\subsection*{Benchmark Datasets}
We assembled three distinct types of data, each providing complementary information to scRNA-seq, for benchmarking HVG selection methods: cell sorting, CITE-seq and single-cell multiome data with paired scRNA-seq and scATAC-seq (MultiomeATAC) (Table~\ref{tab:datatable}). 
Both CITE-seq and single-cell multiome data were generated using 10x Genomics platforms, while cell sorting data were generated using either 10x Genomics, Smart-seq2 \cite{picelli2014full}, or Microwell-seq \cite{han2018mapping}. 

\subsubsection*{Cell Sorting Data}
Each cell sorting dataset contains a mixture of multiple cell types. These datasets are characterized by cells with known cell type labels, determined through cell sorting experiments. These known cell type labels serve as ground truth information for assessing the accuracy of cell clustering based on the gene expression profiles derived from HVGs in scRNA-seq data. 
A total of seven cell sorting datasets have been gathered, featuring varying cell numbers ranging from 3,400 to 131,405  (Table~\ref{tab:datatable}, refer to Availability of data and materials section for references and accession numbers).


\subsubsection*{CITE-seq Data}
Cellular Indexing of Transcriptomes and Epitopes by Sequencing (CITE-seq) generates RNA-seq data alongside measurements of surface proteins in each cell. These surface protein measurements, obtained using antibodies, are termed Antibody-Derived Tags (ADTs).  
Given that RNA-seq and ADTs represent distinct data modalities, ADTs can be utilized to validate results obtained from RNA-seq. A collection of seven CITE-seq datasets was assembled, with cell numbers ranging from 7,665 to 261,729 (Table~\ref{tab:datatable}). 

\subsubsection*{Single-cell Multiome Data (MultiomeATAC)}
The 10x Chromium Single-cell Multiome ATAC + Gene Expression technology enables simultaneous measurement of RNA-seq and ATAC-seq in each cell. ATAC-seq, which stands for Transposase-Accessible Chromatin by sequencing, identifies open chromatin regions in the genome, indicative of active regulatory elements. Given the distinct data modalities represented by scRNA-seq and scATAC-seq, the latter can serve to validate results obtained from scRNA-seq. In addition to the CITE-seq datasets, we compiled five single-cell multiome datasets, with cell numbers ranging from 2,526 to 12,861 (Table~\ref{tab:datatable}).

\subsection*{Data Processing}
\subsubsection*{scRNA-seq}
For the duo4\_pbmc, duo8\_pbmc, duo4un\_pbmc, bmcite and seurat\_cite datasets, processed data after quality control were downloaded and used for this study. For the other datasets, data preprocessing was done using Seurat, with low-quality cells filtered out based on the criteria listed in Additional file 2: Supplementary Table~S1.
The NormalizeData function from Seurat with default setting was used for library size normalization.

For a fair method comparison, the top 2,000 HVGs from each method were used for downstream analyses, consistent with Seurat’s default configuration. With these selected HVGs, a unified procedure was followed to perform the downstream analyses. The raw read counts of HVGs were normalized by library size and scaled by a factor of 10,000. The log-normalized counts were then standardized for each gene to achieve a mean of zero and a standard deviation of one. Subsequently, principal component analysis (PCA) was applied to these standardized expression values. The top 30 principal components (PCs) were used to assess cell similarity. Cell distances were computed as Euclidean distances based on the top 30 PCs. Cells were clustered using the Louvain algorithm (resolution chosen based on the procedure described in the Evaluation Criteria section below). Both clustering and the calculation of within- and between-cluster variances were based on the top 30 PCs.

\subsubsection*{CITE-seq}
For CITE-seq, ADTs were normalized using centered log-ratio transformation (CLR) across cells \cite{hao2021integrated}. The transformed data were then scaled to have zero mean and unit standard deviation. These scaled data were directly used to calculate distances between cells, cluster cells using the Louvain algorithm, and compute within- and between-cluster variances.


\subsubsection*{scATAC-seq}
For single-cell multiome data with paired scRNA-seq and scATAC-seq, the scATAC-seq was processed using the following steps: (1) Filter the data to retain features with nonzero entries in at least 10 cells, and further detailed filtering criteria are discussed in Additional file 2: Supplementary Table~S1; (2) Normalize the data using term frequency–inverse document frequency (TF-IDF); (3) Perform singular value decomposition (SVD) and select the $2^{nd}$ to $30^{th}$ latent semantic indexing (LSI) components; (4) Multiply the LSIs by the corresponding square roots of singular values, without additional scaling, to capture cell-to-cell distances similarly to PCA. These scaled LSIs were also used for cell clustering via Louvain algorithm, and computing within- and between-cluster variances. The first LSI was discarded because of its high correlation with library size (Additional file 1: Supplementary Figure~S3).

\subsection*{Evaluation Criteria}
\subsubsection*{Criteria for Cell Sorting}
In cell sorting datasets, discrete cell type labels were used as orthogonal information to evaluate scRNA-seq analysis results. Accordingly, the following criteria were used for method evaluation:

\begin{itemize}
    \item \textbf{Adjusted rand index (ARI)}: For evaluation using ARI \cite{santos2009use}, two sets of discrete labels were compared. One set consisted of the true cell type labels from cell sorting, while the other set consisted of the clustering labels based on HVGs in scRNA-seq. Louvain clustering was performed at resolutions ranging from 0.1 to 2, increasing in steps of 0.1. Following the approach in \cite{luecken2022benchmarking}, the clustering output with the highest ARI compared to the true labels was used to characterize the performance of a HVG selection method.

    \item \textbf{Normalized mutual information (NMI)}: The input requirements for NMI \cite{lancichinetti2009detecting} are the same as for ARI. NMI is a scaled version of mutual information that quantifies the overlapping information between two sets of cell type labels. Louvain clustering was performed at resolutions ranging from 0.1 to 2, in steps of 0.1. The clustering output with the highest NMI compared to the true labels was used \cite{luecken2022benchmarking}.
    
    \item \textbf{Variance ratio}: The variance ratio is the ratio of between-cell-type variance to within-cell-type variance. Within-cell-type variance is the sum of the total variances of cells within each cell type, characterizing the internal cohesion of cells within the same cell type. Between-cell-type variance is the total variance of all cells minus the within-cell-type variance, characterizing the external separation of different cell types. A higher variance ratio indicates better separation among different cell types.

    \item \textbf{$k$-NN classification accuracy}: HVGs were used to quantify the cell-to-cell distance within the RNA-seq space. This distance metric was used to perform k-nearest neighbors (k-NN) classification ($k=3$). The accuracy of this classification (i.e., percentage of corrected predicted class labels) reflects the effectiveness of the method in distinguishing different cell types based on their RNA-seq profiles. 
    
    \item \textbf{Average silhouette width (ASW)}: The cell-type ASW \cite{rousseeuw1987silhouettes} is the average silhouette width across all cells and cell type pairs, which assesses the balance between a cell's within-cell-type distances and its distances to the closest another cell type. ASW values range from -1 to 1, where -1, 0, and 1 signify significant misclassification, equal variance within and between cell types, and perfect clustering by cell type, respectively. ASW was calculated based on true cell type labels and cell embeddings in the top 30 principal components (PCs) corresponding to HVGs.
    
    \item \textbf{Cell-type local inverse Simpson index (LISI)}: We implemented the refined graph LISI \cite{luecken2022benchmarking} to measure cell type separation, utilizing a $k$-NN integrated graph as the embedded space for distance calculations. LISI scores \cite{korsunsky2019fast} indicate the expected number of neighboring cells to be drawn before encountering a cell of the same type, based on the $k$-NN integrated graph. This graph was constructed using the top 30 PCs corresponding to each set of HVGs, and LISI was calculated using the cell type labels.  LISI values range from 1 to $B$, the number of unique cell types, where 1 indicates perfect separation of cell types and $B$ indicates perfect mixing. For consistent interpretation, we used 1/LISI when ranking HVG selection methods.

\end{itemize}
\subsubsection*{Criteria for CITE-seq and MultiomeATAC data}
In CITE-seq and single-cell multiome datasets, protein levels measured by ADTs and chromatin accessibility measured by scATAC-seq were  utilized as orthogonal information to evaluate scRNA-seq analysis results. Both ADTs and LSI embeddings of scATAC-seq yield continuous values. Therefore, the following criteria were used for method evaluation:
\begin{itemize}
    \item \textbf{Variance ratio}:  Cells are clustered using Louvain clustering on HVGs derived from scRNA-seq data. The variance ratio is calculated as the ratio of between-cell-cluster variance to within-cluster variance. However, this variance is computed in an orthogonal information space, specifically in the space defined by ADTs of CITE-seq or LSI embeddings of scATAC-seq data. This approach is analogous to the variance ratio used for cell sorting, with a key difference being that cell types identified through cell sorting are replaced by cell clusters identified through scRNA-seq. Additionally, the variance is calculated using CITE-seq ADTs or scATAC-seq LSIs instead of scRNA-seq data. A higher variance ratio indicates better separation between different cell clusters, suggesting that the clusters are more distinct in the orthogonal information space. To ensure consistency and comparability, the resolution parameter for Louvain clustering is fixed at $0.1$ during the computation of the variance ratio. This prevents the introduction of more clusters solely to increase the variance ratio since more clusters will lead to a higher variance ratio. 

    \item \textbf{$k$-NN regression mean square error (MSE)}: The $k$-NN regression MSE is analogous to $k$-NN classification accuracy but applied to continuous data. The RNA-seq space is used to identify the nearest neighbors, with distances computed based on HVGs. The $k$-NN MSE is calculated using orthogonal data, such as CITE-seq ADTs or scATAC-seq LSIs ($k=3$). MSE measures the average squared differences between the predicted values (based on $k$-NN regression) and the actual values in the orthogonal information space. It reflects how well the nearest neighbors in the RNA-seq space correspond to relationships in the orthogonal data. To facilitate interpretation, we use 1/MSE when ranking HVG selection methods. A higher 1/MSE value indicates better performance, meaning that the nearest neighbors in the RNA-seq space are more accurately reflected in the orthogonal data space.

    \item \textbf{$k$-NN distance ratio}: The $k$-NN distance ratio is the ratio of the distance to the cells outside a cell's $k$-nearest neighbors to the distance to the $k$-nearest neighbors ($k=100$). Here nearest neighbors are determined using scRNA-seq, and the distance ratio is calculated using orthogonal data, such as CITE-seq ADTs or scATAC-seq LSIs. The distance is computed as the sum of distances between each pair of cells. 
    A higher distance ratio indicates better separation of different cell types, as it shows that cells are closer to their own cluster members than to members of other clusters.
    
    \item \textbf{Distance correlation}:  Distance correlation is used to assess the consistency of cell-to-cell distances across different data types. Distances are computed between each pair of cells within the dataset. These distances can be calculated in two spaces: (1) RNA-seq space, defined by HVGs; (2) orthogonal information space, defined by CITE-seq ADTs or scATAC-seq LSIs. The Pearson correlation coefficient is used to compute the correlation between the distance lists from the two different spaces.
    
    \item \textbf{Adjusted rand index (ARI)}: ARI is computed using  two sets of clustering labels. One set is derived from the RNA-seq space using HVGs. The other set is derived from the CITE-seq ADT or scATAC-seq LSI space. Louvain clustering is performed in both spaces. The clustering is done across a range of resolutions from 0.1 to 2 in steps of 0.1 for principal components derived from HVGs in the scRNA-seq space, ADTs, and scATAC-seq LSIs, respectively. ARI is computed for each pair of clustering results (one from RNA-seq space, one from ADT or scATAC-seq space). The highest ARI value across all pairs is used as the final output.
    A higher ARI indicates better alignment between clusters derived from RNA-seq data and those derived from ADT or scATAC-seq data, implying more consistent biological clustering.

    \item \textbf{Normalized mutual information}: NMI is another metric used to evaluate the similarity between two clustering results. Similar to ARI, NMI requires two sets of clustering labels, one set from the RNA-seq space using HVGs and the other set from the CITE-seq ADT or scATAC-seq LSI space. Louvain clustering is performed on both datasets, with resolutions ranging from 0.1 to 2 in steps of 0.1 for PCs derived from HVGs, ADTs, and scATAC-seq LSIs. NMI is calculated for each pair of clustering results (one from RNA-seq space, one from ADT or scATAC-seq space). The highest NMI value from these computations is used as the final output. Higher NMI values indicate greater similarity and consistency between the clusters identified in different data modalities, reflecting better overall clustering performance.
\end{itemize}

\section*{Additional Files}
\begin{itemize}
    \item Additional file 1: Supplementary Figures~S1-S6
    \begin{itemize}
    \item Figure~S1 --- A comparison of seven existing methods and mixHVG (default) in the zheng\_pbmc dataset.
    \item Figure~S2 --- A comparison of seven existing methods and mixHVG (default) in the cbmc8k\_cite dataset.
    \item Figure~S3 --- Pearson correlation between cell's sequencing depth for scATAC (i.e., library size) and the first 10 LSI components in the single-cell multiome datasets.
    \item Figure~S4 --- A comparison of seven existing methods and mixHVG (default) in the pbmc10k\_multi dataset.
    \item Figure~S5 --- Comparison of 21 baseline HVG selection methods across all benchmark datasets and criteria from the view of data transformation.
    \item Figure~S6 --- Heatmap showing the number of overlapping genes selected by different HVG methods.
    \end{itemize}
    \item Additional file 2: Supplementary Table~S1 --- Criteria for filtering low quality cells in scRNA-seq.
\end{itemize}
    
\section*{Acknowledgements}%% if any
The authors would like to thank Wentao Zhan from the Ji Lab for helpful discussions, and Dr. Zhicheng Ji from Duke University for sharing some processed datasets. 

\section*{Funding}%% if any
This research is supported by The National Institutes of Health grants R01HG010889 and R01HG009518 to H.J.

\section*{Abbreviations}%% if any
\begin{itemize}
    \item {\bf ADT}: Antibody-Derived Tags
    \item {\bf ARI}: Adjusted Rand Index
    \item {\bf ASW}: Average Silhouette Width
    \item {\bf CITE-seq}: Cellular Indexing of Transcriptomes and Epitopes by Sequencing
    \item {\bf HVG}: Highly Variable Gene
    \item {\bf LISI}: Local Inverse Simpson Index
    \item {\bf LOESS}: Locally weighted (or estimated) scatterplot  smoother
    \item {\bf LSI}: Latent Semantic Indexing
    \item {\bf NMI}: Normalized Mutual Information
    \item {\bf PBMC}: Peripheral Blood Mononuclear Cell
    \item {\bf PC}: Principal Component
    \item {\bf PCA}: Principal Component Analysis
    \item {\bf scATAC-seq}: Single-cell Assay for Transposase-Accessible Chromatin using sequencing
    \item {\bf scRNA-seq}: Single-cell RNA sequencing
    %\item {\bf NGS}: Next Generation Sequencing
    \item {\bf SCT}: sctransform
    \item {\bf SVD}: Singular Value Decomposition
    \item {\bf TFIDF}: Term Frequency–Inverse Document Frequency
\end{itemize}


\section*{Availability of data and materials}
The data used in this analysis are all publicly available. All data are described in the Table~\ref{tab:datatable}. Cell sorting: GBM\_sd (GSE84465) \cite{darmanis2017single}, duo4\_pbmc \cite{duo2018systematic}, duo8\_pbmc \cite{duo2018systematic}, duo4un\_pbmc \cite{duo2018systematic}, zheng\_pbmc \cite{zheng2017massively}, mus\_tissue \cite{han2018mapping}, homo\_tissue \cite{han2018mapping}; %mus\_tissue\_large \cite{han2018mapping};
CITE-seq: pbmc\_cite (GSE100866) \cite{stoeckius2017large,linderman2022zero}, cbmc8k\_cite (GSE100866) \cite{stoeckius2017large,linderman2022zero},  FBM\_cite (GSE166895) \cite{jardine2021blood}, FLiver\_cite (GSE166895) \cite{jardine2021blood}, bmcite (GSE128639) \cite{stuart2019comprehensive}, seurat\_cite \cite{hao2021integrated}, sucovid\_cite \cite{su2020multi}; Single-cell multiome data with paired scRNA-seq and scATAC-seq: all datasets go for section(Single-cell Multiome ATAC + Gene Expression) of (\url{https://www.10xgenomics.com/resources/datasets}): pbmc3k\_multi, homo\_brain3k\_multi, mus\_brain5k\_multi, pbmc10k\_multi, lymphoma\_multi. The preprocessing code is available under \url{https://github.com/RuzhangZhao/scRNAseqProcess}, and the processed data is available under \url{https://zenodo.org/records/12135988}. Detailed downloading links are also included. Codes for the benchmark analysis can be found under \url{https://github.com/RuzhangZhao/BenchmarkHVG}.

\section*{Availability of software}
The R package \texttt{mixhvg} is available at \url{https://CRAN.R-project.org/package=mixhvg} \cite{mixmix2024}. The R package \texttt{benchmarkHVG} is available at \url{https://github.com/RuzhangZhao/benchmarkHVG}. 

\section*{Ethics approval and consent to participate}%% if any
Not applicable.

\section*{Competing interests}
The authors declare that they have no competing interests.


\section*{Consent for publication}%% if any
Not applicable.


\section*{Authors' contributions}
RZ, WZ and HJ conceived the study. RZ developed software and benchmark pipeline. RZ and JL performed the data analyses. HJ supervised the study. RZ and HJ wrote the initial manuscript. JL, WZ and NZ reviewed the manuscript and provided feedback. All authors approved the final manuscript.
\bibliography{sn-article}% common bib file

\renewcommand{\thefigure}{S1}
\begin{figure}[h!]
    \centering
    \includegraphics[width=\textwidth]{Additional_File1_FigureS1.png}
    \caption{A comparison of seven existing methods and mixHVG (default) in the zheng\_pbmc dataset. Each subplot depicts a UMAP generated using scRNA-seq data and HVGs selected by a specific method.  The color indicates true cell type labels obtained from cell sorting. The plot includes seven publicly available methods denoted by an asterisk (*) and the default hybrid method of mixHVG (mv\_lognc\_scran(1mv) + poisson\_scran(3pos) + disp\_nc\_seuratv1(4dis)). The evaluation criteria values are displayed for each method. Cell display orders are shuffled to prevent overlap between different cell types.}
    \label{fig:zheng}
\end{figure}

\renewcommand{\thefigure}{S2}
\begin{figure}[h!]
    \centering
    \includegraphics[width=\textwidth]{Additional_File1_FigureS2.png}
    \caption{A comparison of seven existing methods and mixHVG (default) in the cbmc8k\_cite dataset. The cells are colored by the clustering results using ADT with Louvain clustering (resolution 0.2). 
    The plot includes 7 publicly available methods denoted by an asterisk (*) and the default hybrid method of mixHVG (mv\_lognc\_scran(1mv) + poisson\_scran(3pos) + disp\_nc\_seuratv1(4dis)). The values of all criteria are marked for each method. The cells are all shuffled to avoid the case where one cell type are totally covered by the other.}
    \label{fig:cbmc8k}
\end{figure}

\renewcommand{\thefigure}{S3}
\begin{figure}[h!]
    \centering
    \includegraphics[width=\textwidth]{Additional_File1_FigureS3.pdf}
    \caption{Pearson correlation between cell's sequencing depth for scATAC (i.e., library size) and the first 10 LSI components in the single-cell multiome datasets.
    }
    \label{fig:lsi}
\end{figure}

\renewcommand{\thefigure}{S4}
\begin{figure}[h!]
    \centering
    \includegraphics[width=\textwidth]{Additional_File1_FigureS4.png}
    \caption{A comparison of seven existing methods and mixHVG (default) in the pbmc10k\_multi dataset.
    The cells are colored by the clustering results using the $2^{\text{nd}}$ to $30^{\text{th}}$ dimensional LSIs with Louvain clustering (resolution 0.2). The plot includes 7 publicly available methods denoted by an asterisk (*) and the default hybrid method of mixHVG (mv\_lognc\_scran(1mv) + poisson\_scran(3pos) + disp\_nc\_seuratv1(4dis)). The values of all criteria are marked for each method. The cells are all shuffled to avoid the case where one cell type are totally covered by the other.}
    \label{fig:lym}
\end{figure}


\renewcommand{\thefigure}{S5}
\begin{figure}[h!]
    \centering
    \includegraphics[width=\textwidth]{Additional_File1_FigureS5.pdf}
    \caption{Comparison of 21 baseline HVG selection methods across all benchmark datasets and criteria from the view of data transformation. The methods are categorized into four main groups based on input data transformation type: count, normalized count, log-normalized count, and PFlogPF. Each category is represented by a different color. The plot is a reordering of Figure~5. 
    }
    \label{fig:reorder}
\end{figure}

\renewcommand{\thefigure}{S6}
\begin{figure}[h!]
    \centering
    \includegraphics[width=\textwidth]{Additional_File1_FigureS6.pdf}
    \caption{Heatmap showing the number of overlapping genes selected by different HVG methods. Each subplot represents one dataset. Red indicates high overlap.}
    \label{fig:overlap2}
\end{figure}

\renewcommand{\thetable}{S1}
\begin{table}[h!]
    \centering
    \caption{Criteria for filtering low quality cells in scRNA-seq.}
\scriptsize{
    \begin{tabular}{cc}%{|c|c|c|c|c|}
\toprule
\textbf{Dataset} & \textbf{Filtering criteria}  \\
\midrule
GBM\_sd & Keep cells with $8\times 10^{4}\leq$ nCount\_RNA $\leq 9\times 10^{5}$  \\
& \\
mus\_tissue & $\begin{array}{c}\text {Keep cells with nFeature\_RNA $\geq 1000$ } \\ \text{and genes expressed in $\geq$  0.1\% of cells}\end{array}$ \\
&\\
zheng\_pbmc & $\begin{array}{c}\text {Keep cells with nFeature\_RNA $\geq 1000$ } \\ \text{and genes expressed in $\geq$  0.1\% of cells}\end{array}$ \\
& \\
homo\_tissue & $\begin{array}{c}\text {Keep cells with nFeature\_RNA $\geq 500$ } \\ \text{and genes expressed in $\geq$  0.1\% of cells}\end{array}$\\
& \\
pbmc\_cite& Remove genes and cells associated with mouse\\
& \\
cbmc8k\_cite& Remove genes and cells associated with mouse\\
& \\
FLiver\_cite & Keep cells with both scRNA-seq and ADT\\
& \\
FBM\_cite& Keep cells with both scRNA-seq and ADT\\
& \\
sucovid\_cite & Follow the processing protocol in \cite{zhang2021tree}\\
& \\
pbmc3k\_multi & $\begin{array}{c}\text {Keep cells with $1\times 10^{3} \leq$ nCount\_ATAC $\leq 5\times 10^{4}$} \\ \text{$1\times 10^{3} \leq$ nCount\_RNA $\leq 2.5\times 10^{4}$, percent.mt $\leq 20$}\end{array}$\\
& \\
homo\_brain3k\_multi & $\begin{array}{c}\text {Keep cells with $1\times 10^{3} \leq$ nCount\_ATAC $\leq 2\times 10^{5}$} \\ \text{$8\times 10^{2} \leq$ nCount\_RNA $\leq 5\times 10^{4}$, percent.mt $\leq 3$}\end{array}$\\
& \\
mus\_brain5k\_multi& $\begin{array}{c}\text {Keep cells with $1\times 10^{3} \leq$ nCount\_ATAC  $\leq 1\times 10^{5}$} \\ \text{$1\times 10^{3} \leq$ Count\_RNA $\leq 5\times 10^{4}$}\end{array}$\\
& \\
pbmc10k\_multi& $\begin{array}{c}\text{Keep cells with $5\times 10^{3} \leq$ nCount\_ATAC  $\leq 7\times 10^{4}$} \\ \text{$1\times 10^{3} \leq$ nCount\_RNA $\leq 2.5\times 10^{4}$, percent.mt $\leq 20$}\end{array}$\\
& \\
lymphoma\_multi&$\begin{array}{c}\text {Keep cells with $5\times 10^{2} \leq$  nCount\_ATAC $\leq 7\times 10^{4}$} \\ \text{$3\times 10^{2} \leq$ nCount\_RNA $\leq 2\times 10^{4}$, percent.mt $\leq 3$}\end{array}$\\
\botrule
\end{tabular}
}

\raggedright{Note: nFeature\_RNA: number of genes detected per cell in scRNA-seq. nCount\_RNA: number of UMIs per cell in scRNA-seq. percent.mt: mitochondrial percentage of a cell in scRNA-seq. nCount\_ATAC: number of sequencing reads detected per cell in scATAC-seq.}
\label{tab:preprocess}
\end{table}


%% if required, the content of .bbl file can be included here once bbl is generated
%%\input sn-article.bbl


\end{document}
