## Supplementary figures and images for "A systematic evaluation of highly variable gene selection methods for single-cell RNA-sequencing"

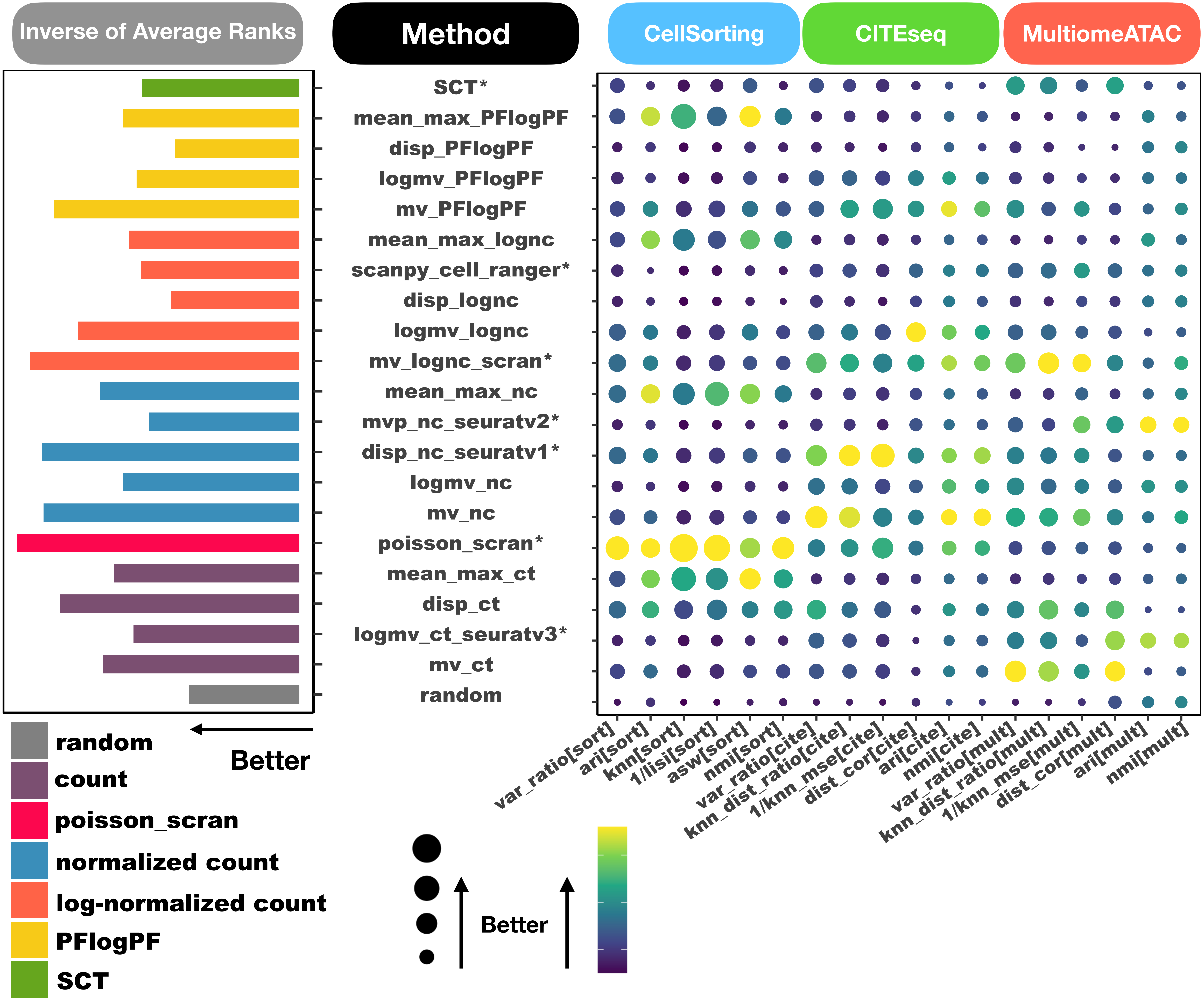

Supplement: The full latex folder for manuscript [file 608519_file02.zip › [submission]HVG_sn_for_biorXiv/Additional_File1_FigureS5.pdf]

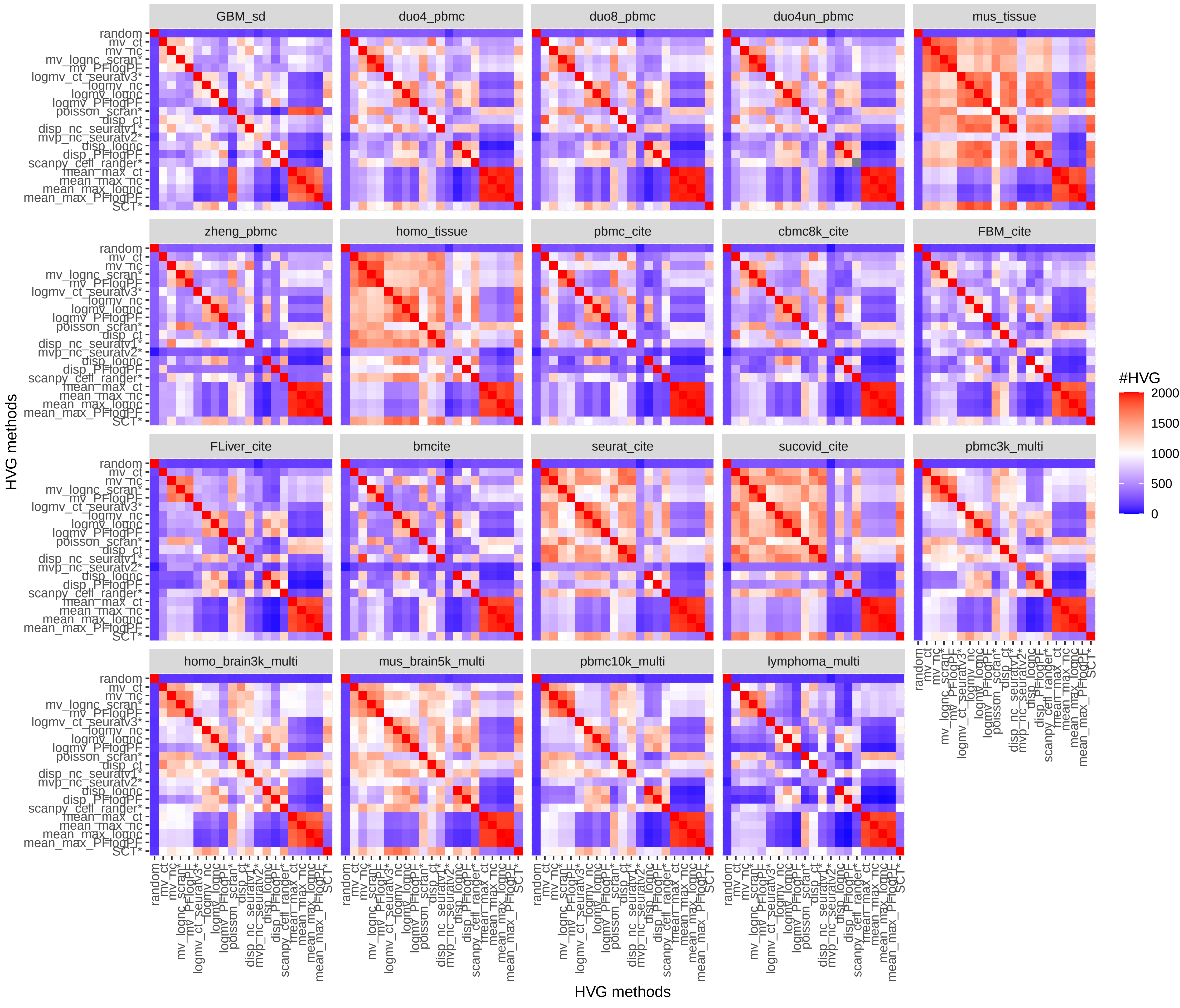

Supplement: The full latex folder for manuscript [file 608519_file02.zip › [submission]HVG_sn_for_biorXiv/Additional_File1_FigureS6.pdf]

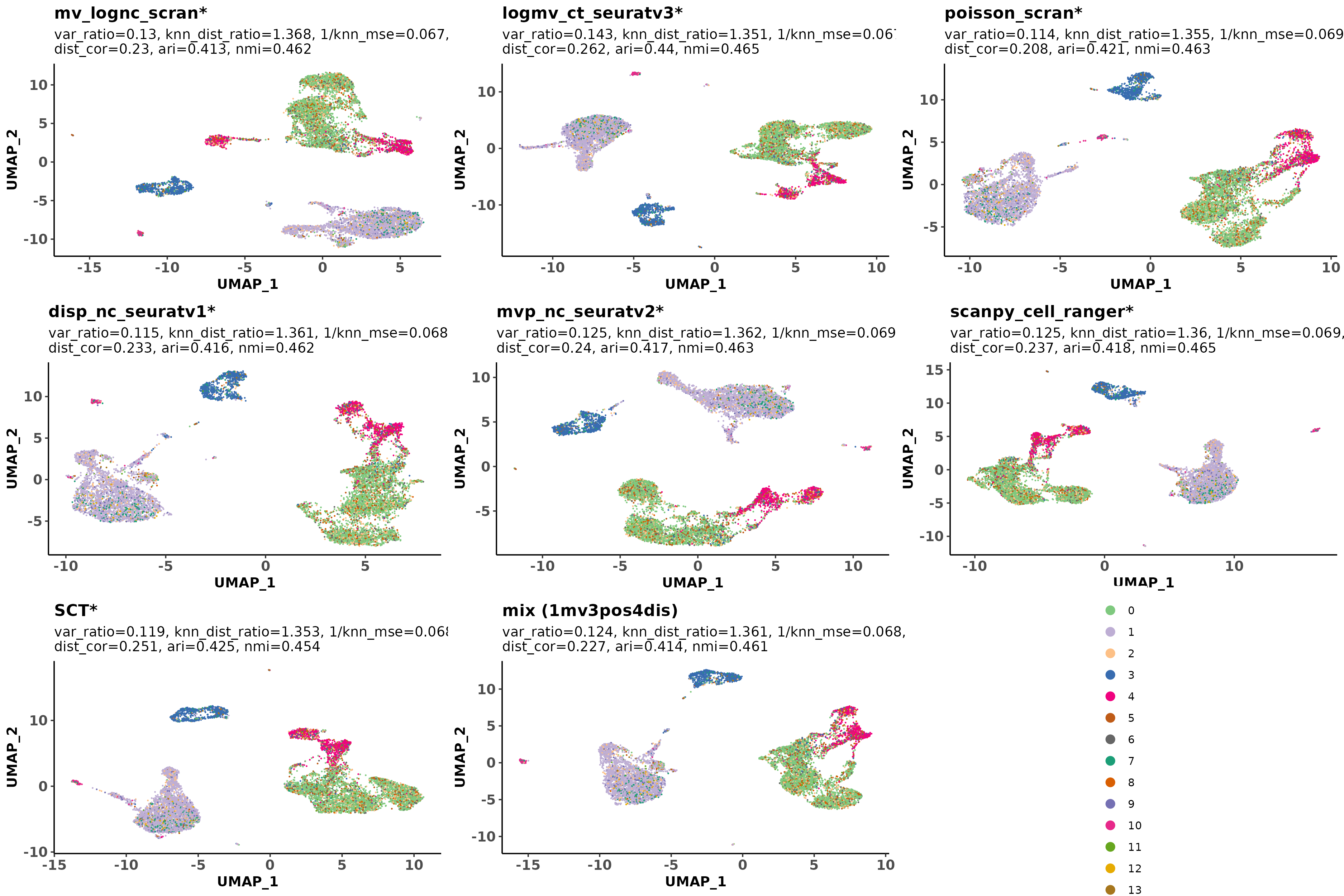

Supplement: The full latex folder for manuscript [file 608519_file02.zip › [submission]HVG_sn_for_biorXiv/Additional_File1_FigureS4.png]

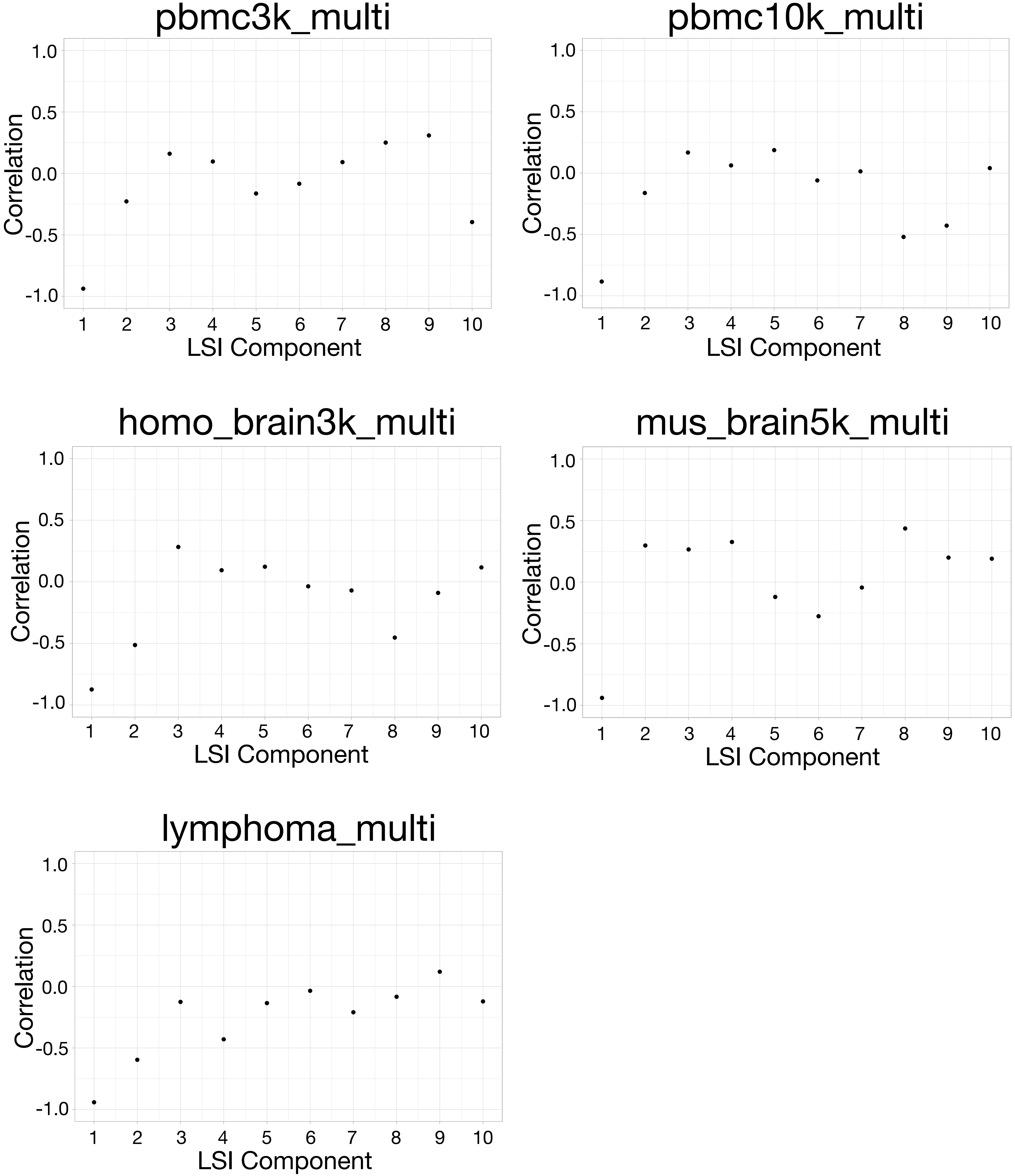

Supplement: The full latex folder for manuscript [file 608519_file02.zip › [submission]HVG_sn_for_biorXiv/Additional_File1_FigureS3.pdf]

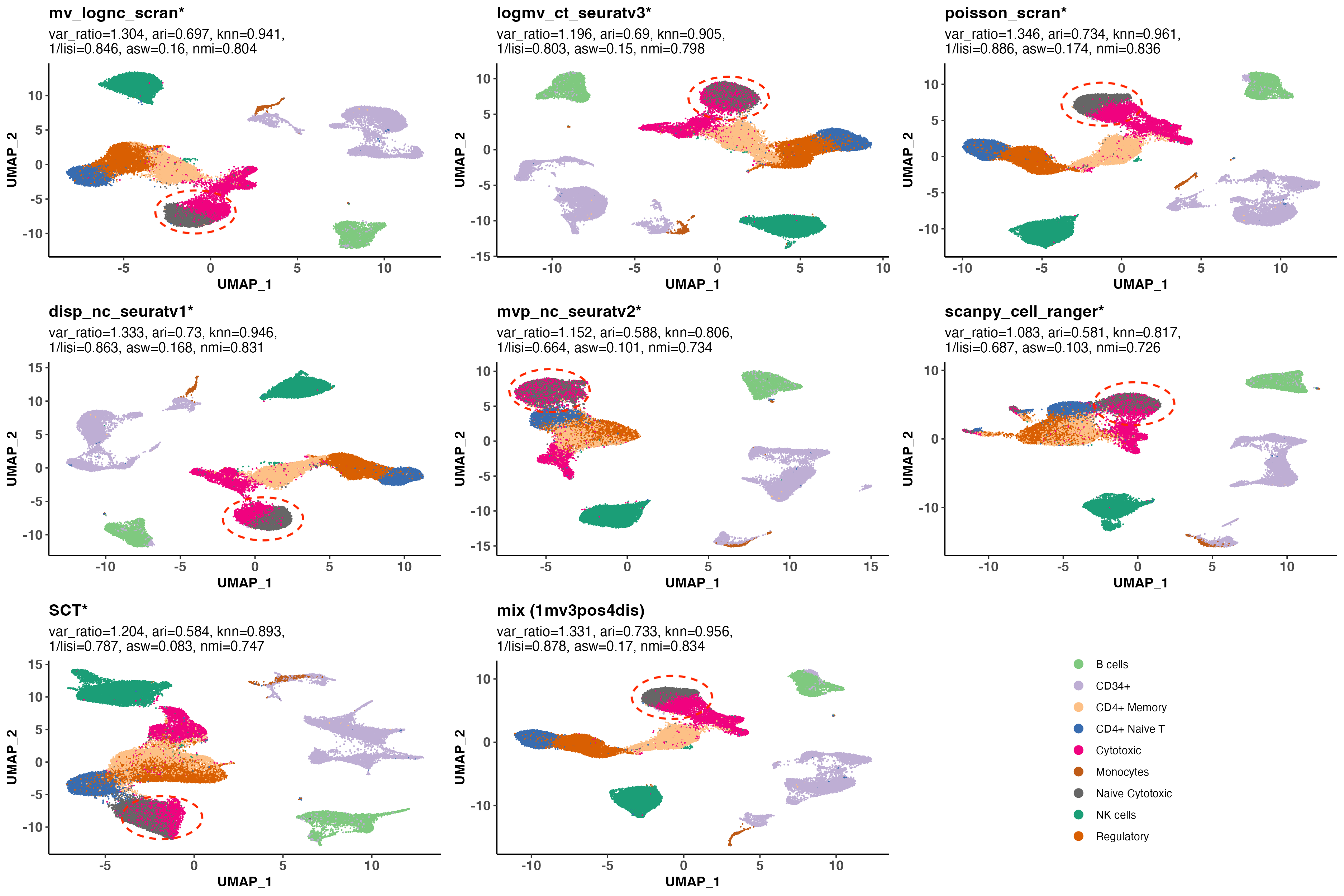

Supplement: The full latex folder for manuscript [file 608519_file02.zip › [submission]HVG_sn_for_biorXiv/Additional_File1_FigureS1.png]

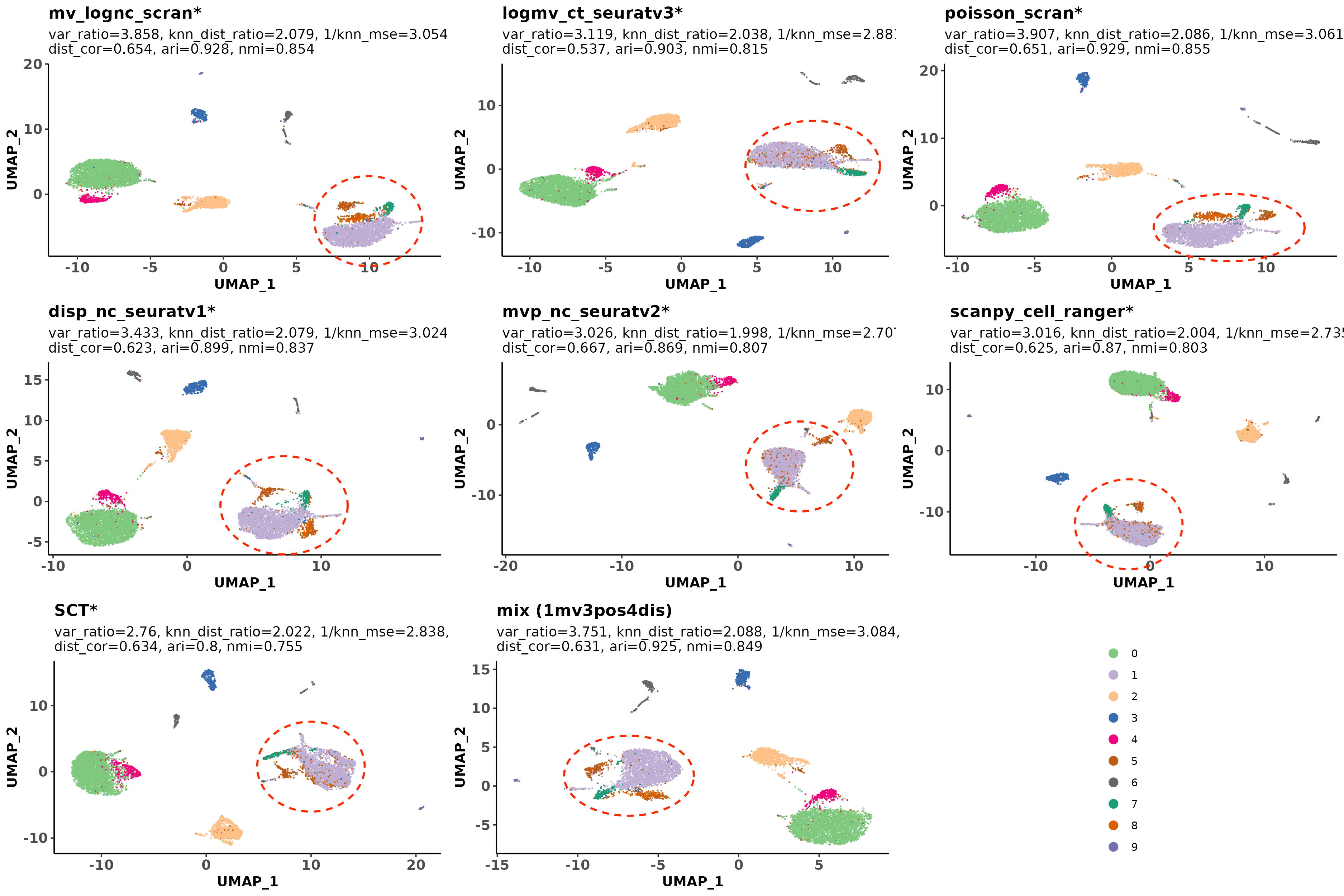

Supplement: The full latex folder for manuscript [file 608519_file02.zip › [submission]HVG_sn_for_biorXiv/Additional_File1_FigureS2.png]

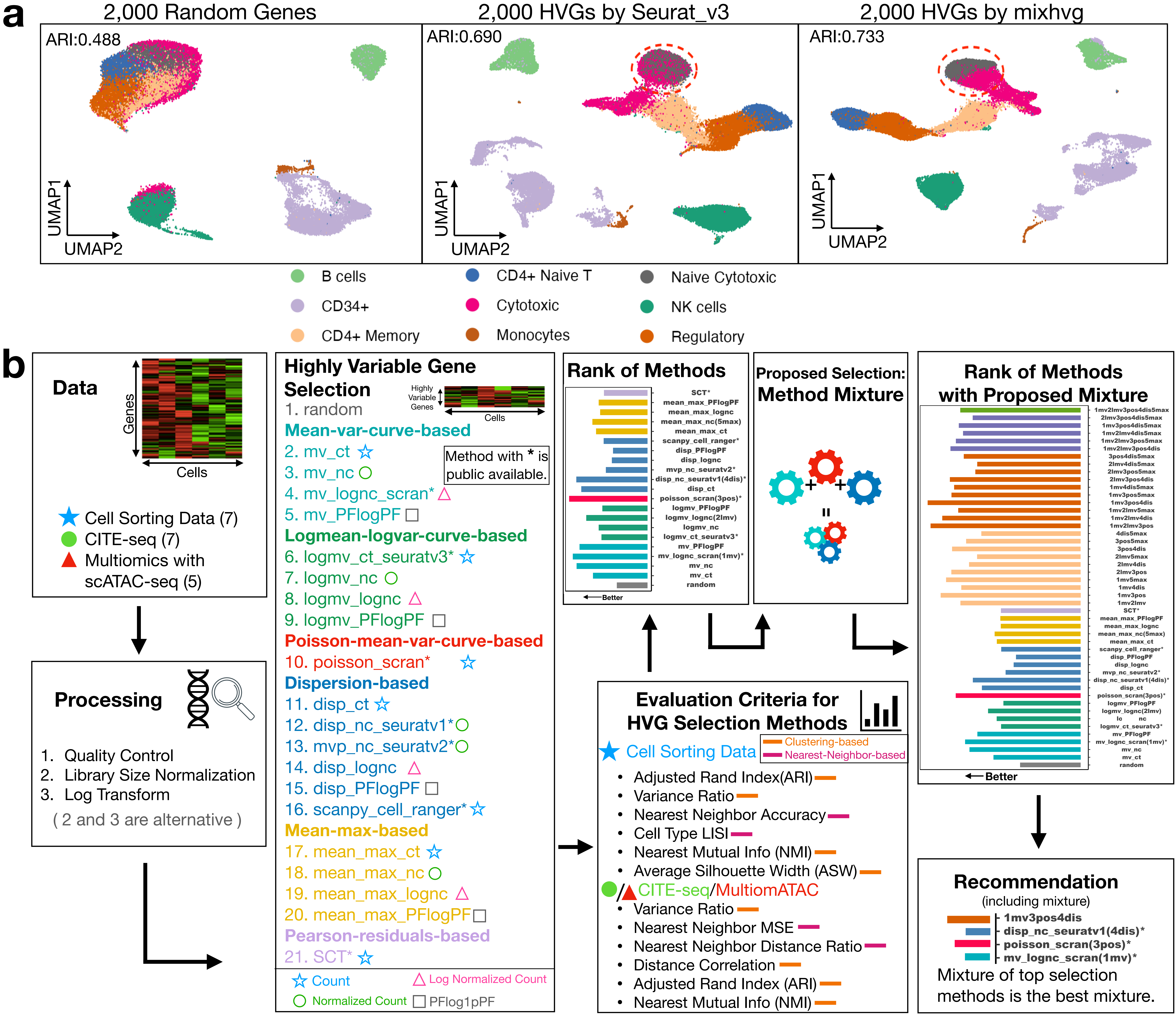

Supplement: The full latex folder for manuscript [file 608519_file02.zip › [submission]HVG_sn_for_biorXiv/Figure1.pdf]

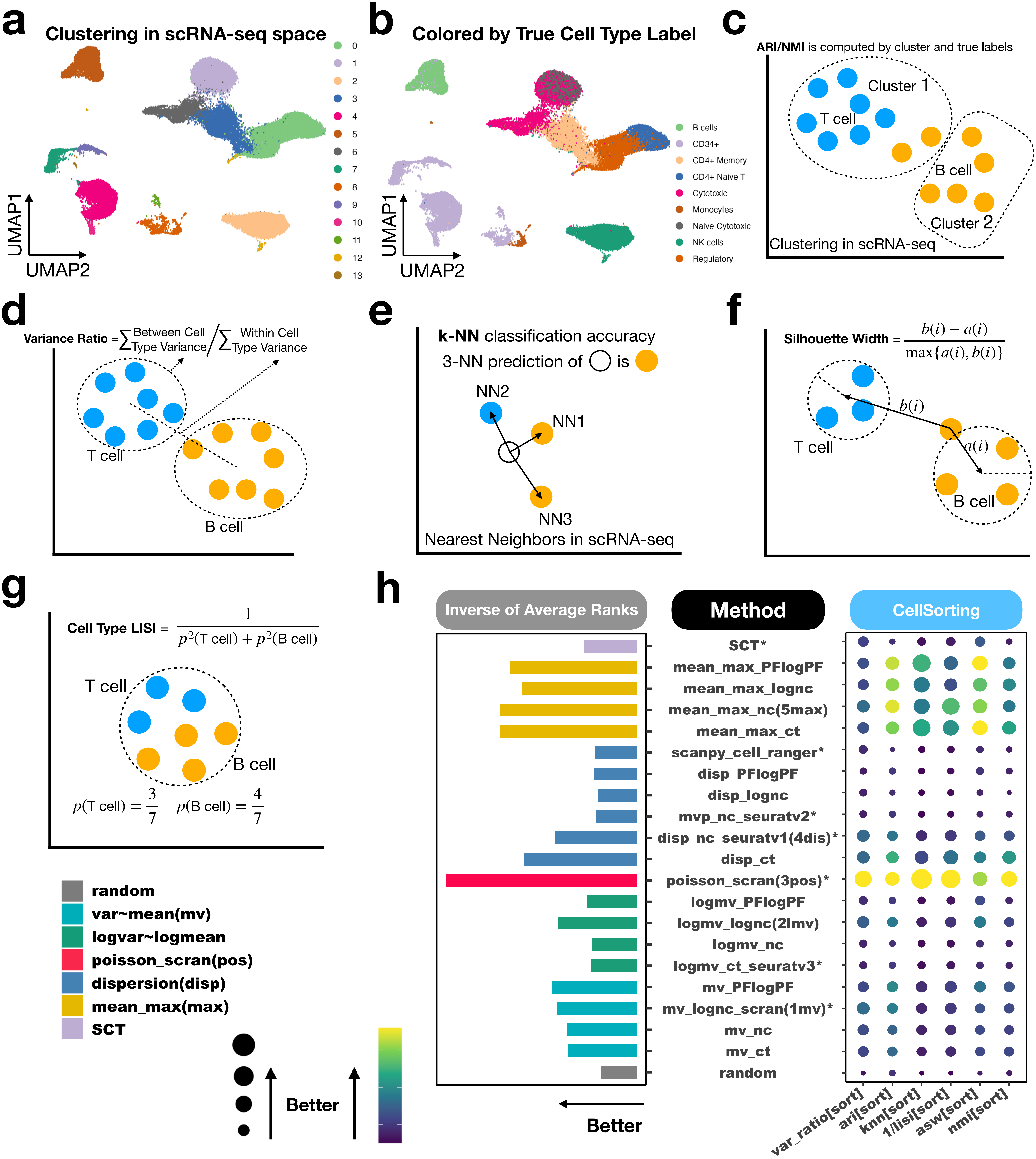

Supplement: The full latex folder for manuscript [file 608519_file02.zip › [submission]HVG_sn_for_biorXiv/Figure2.pdf]

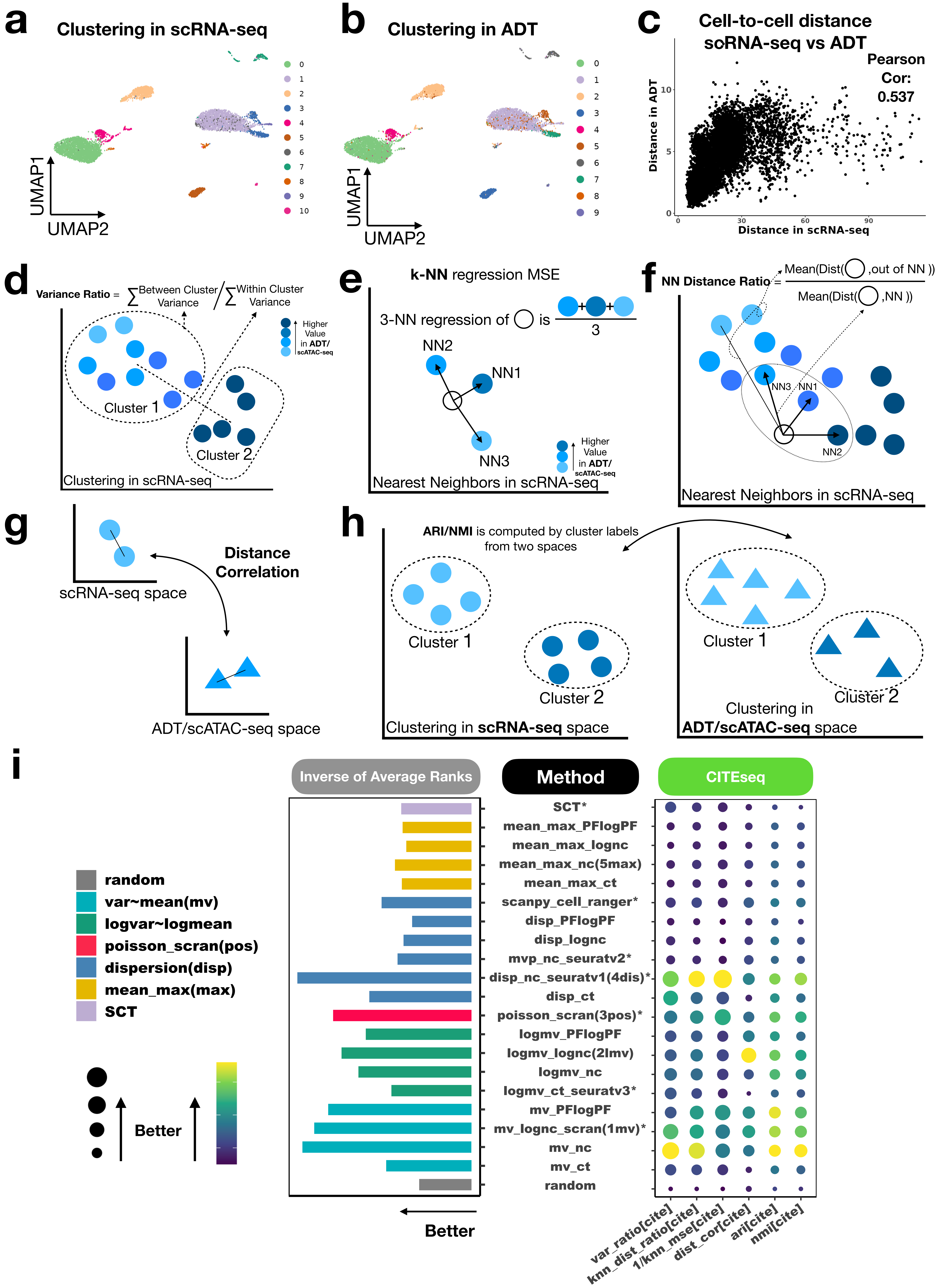

Supplement: The full latex folder for manuscript [file 608519_file02.zip › [submission]HVG_sn_for_biorXiv/Figure3.pdf]

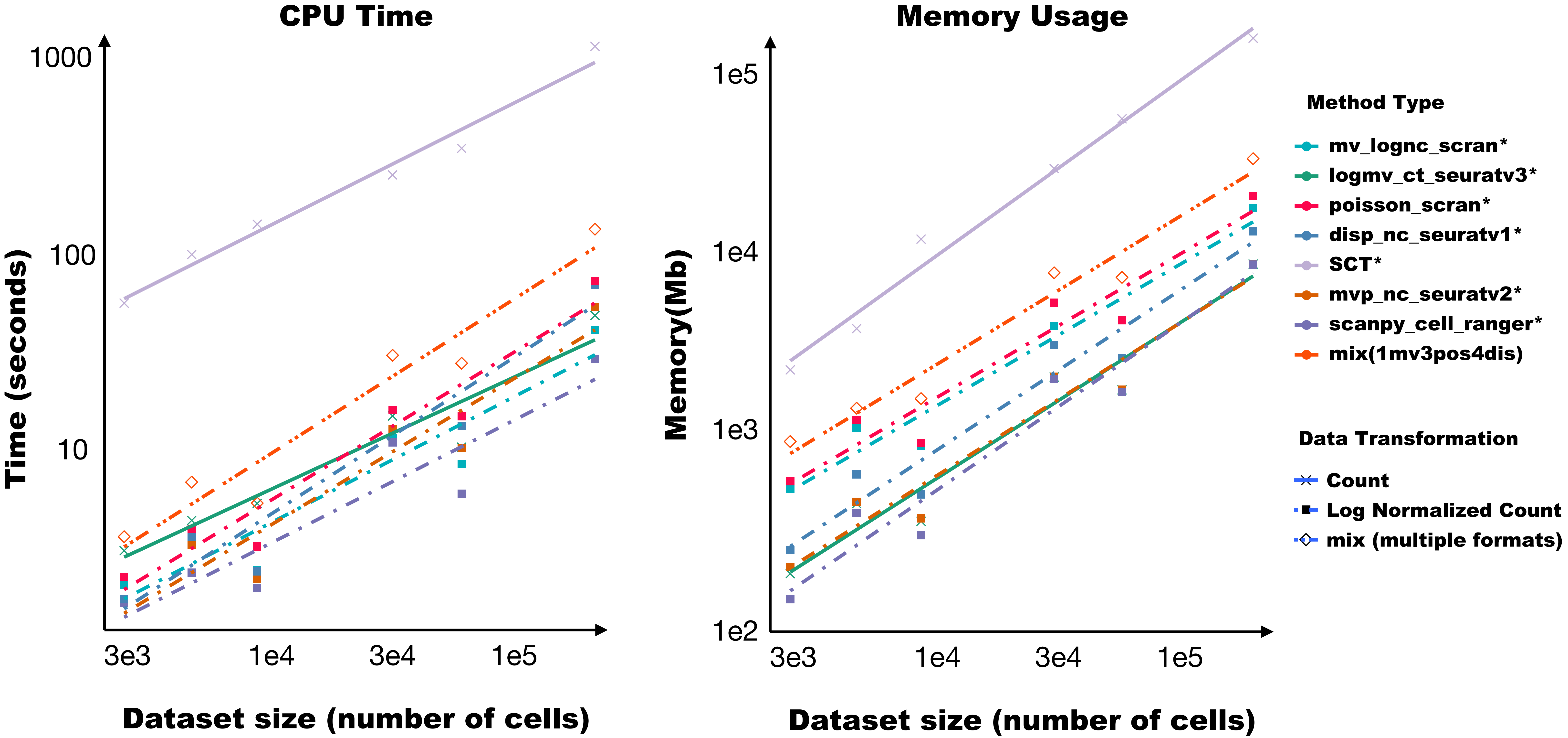

Supplement: The full latex folder for manuscript [file 608519_file02.zip › [submission]HVG_sn_for_biorXiv/Figure7.pdf]

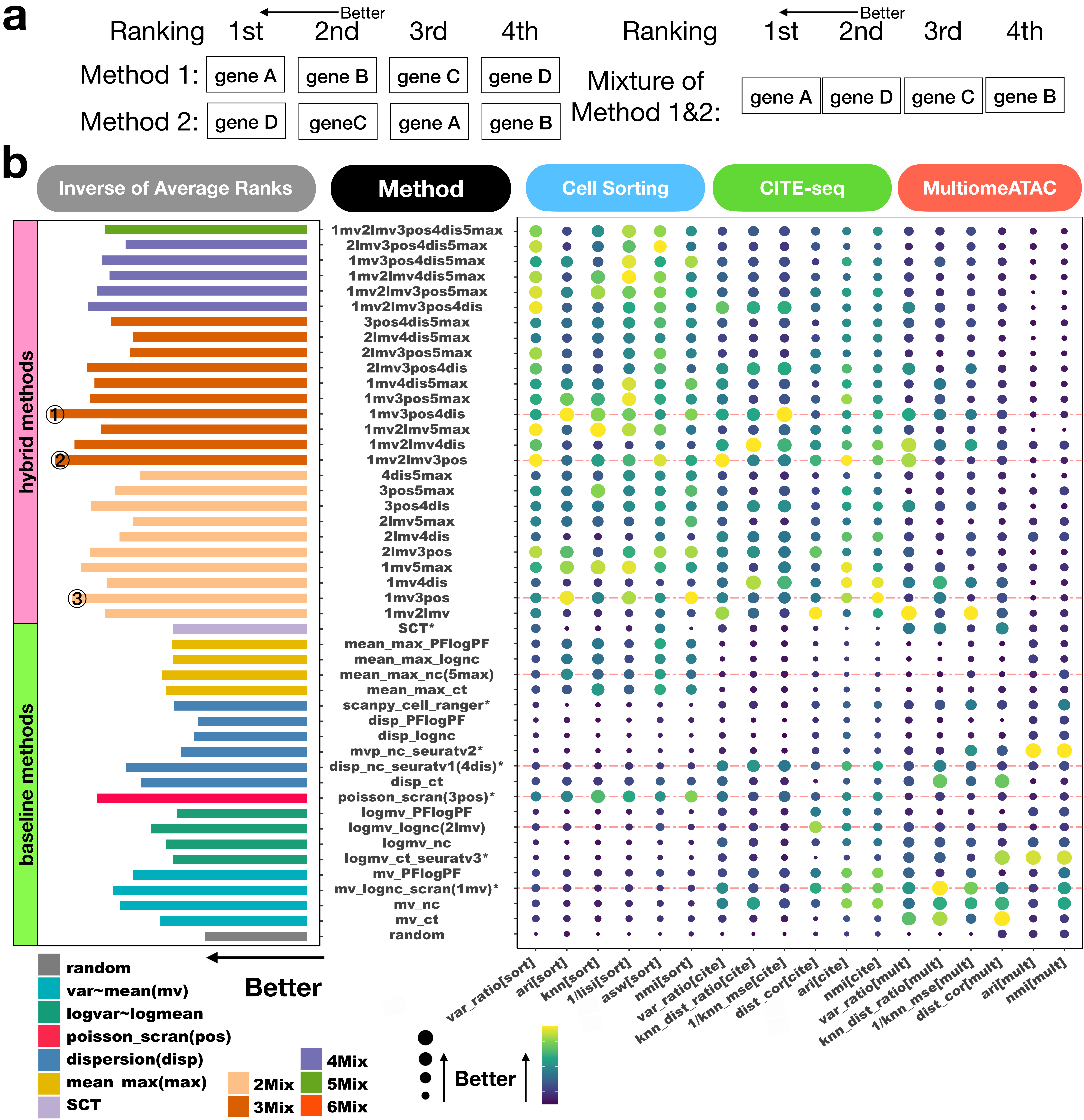

Supplement: The full latex folder for manuscript [file 608519_file02.zip › [submission]HVG_sn_for_biorXiv/Figure6.pdf]

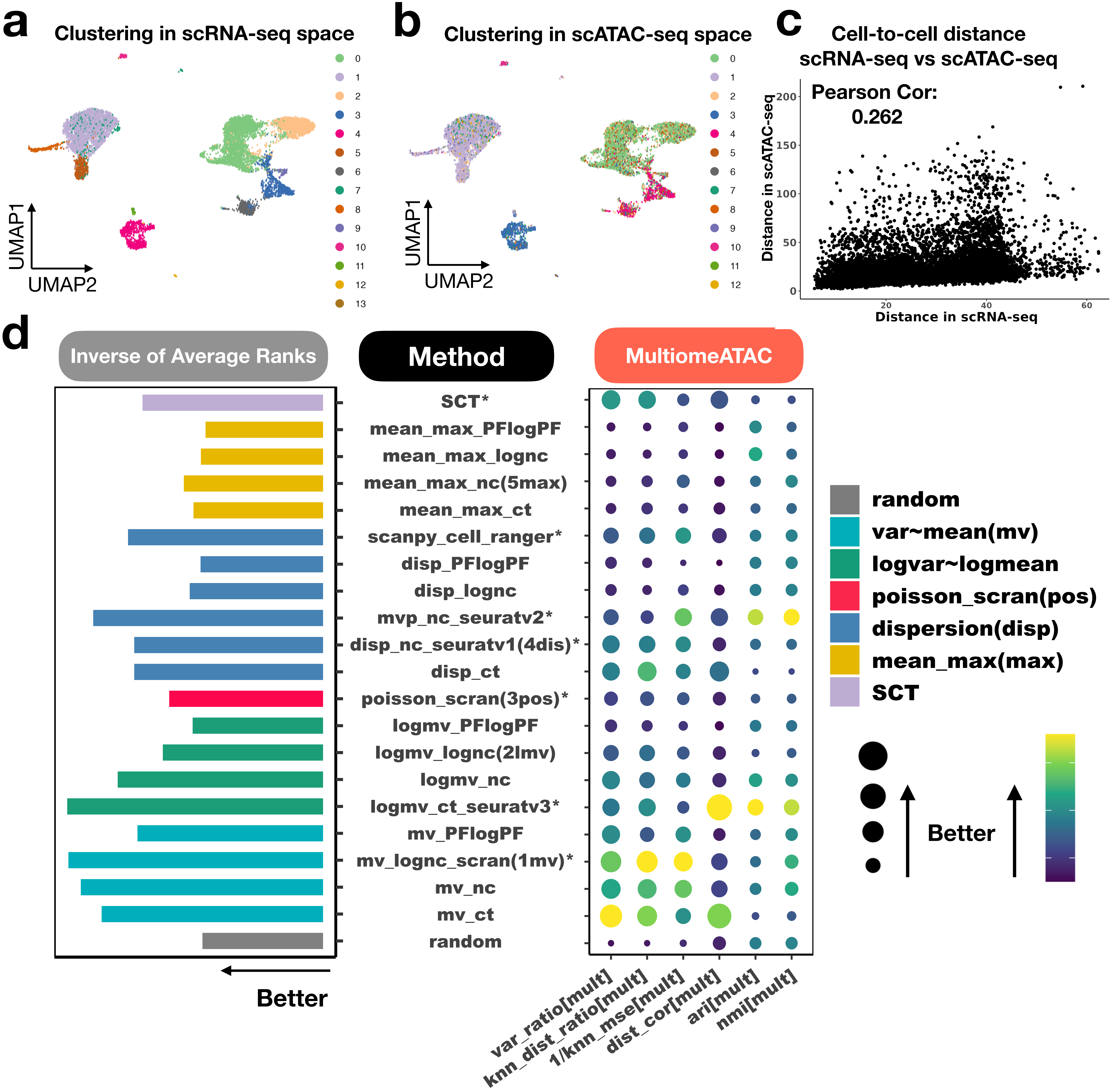

Supplement: The full latex folder for manuscript [file 608519_file02.zip › [submission]HVG_sn_for_biorXiv/Figure4.pdf]

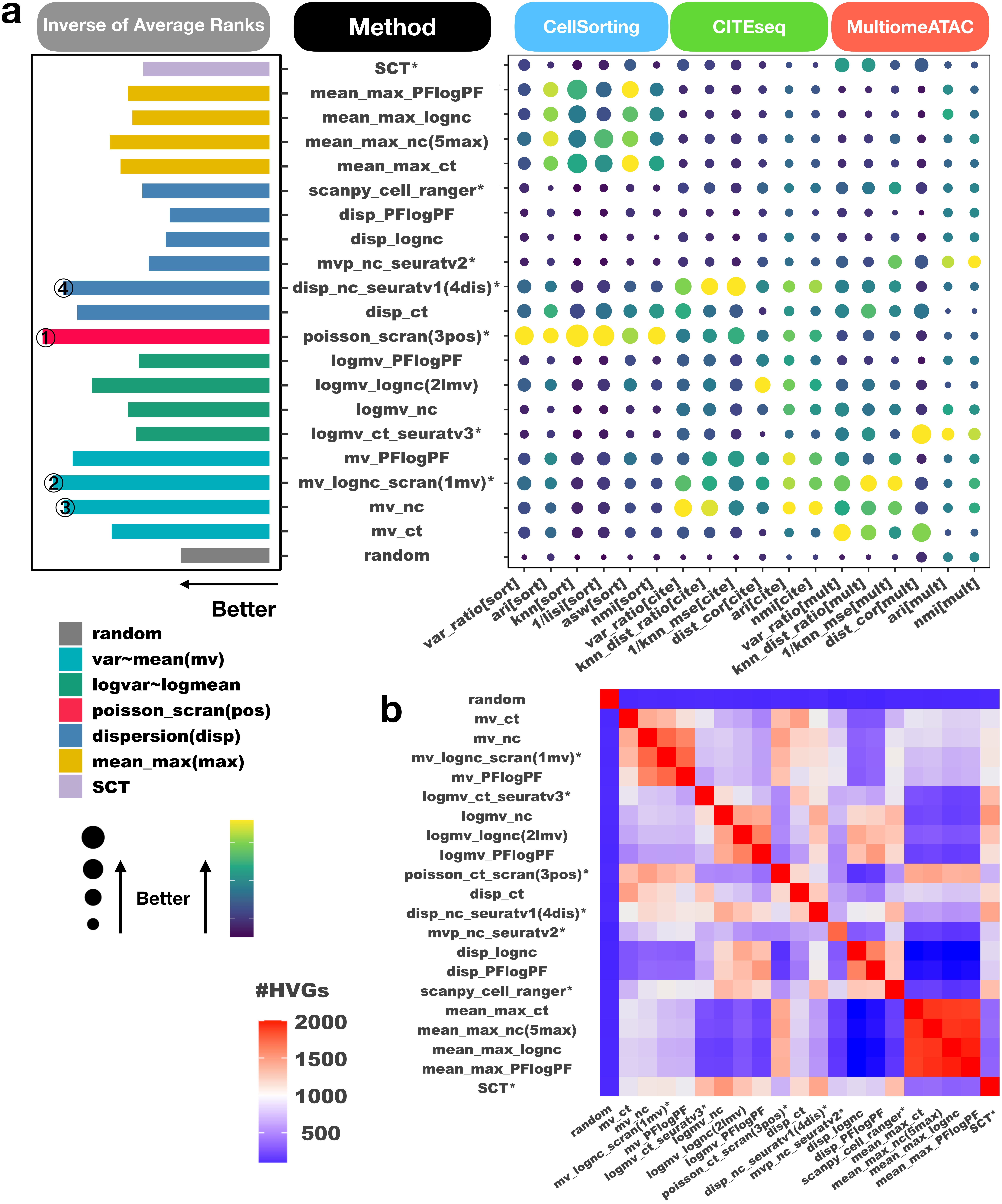

Supplement: The full latex folder for manuscript [file 608519_file02.zip › [submission]HVG_sn_for_biorXiv/Figure5.pdf]
